# PADI2-mediated citrullination is required for efficient oligodendrocyte differentiation and myelination

**DOI:** 10.1101/425348

**Authors:** Ana Mendanha Falcão, Mandy Meijer, Antonella Scaglione, Puneet Rinwa, Eneritz Agirre, Jialiang Liang, Sara C. Larsen, Abeer Heskol, Rebecca Frawley, Michael Klingener, Manuel Varas-Godoy, Alexandre A.S.F. Raposo, Patrik Ernfors, Diogo S. Castro, Michael L. Nielsen, Patrizia Casaccia, Gonçalo Castelo-Branco

**Affiliations:** Laboratory of Molecular Neurobiology, Department of Medical Biochemistry and Biophysics, Karolinska Institutet, Stockholm, Sweden; Icahn School of Medicine at Mount Sinai, New York, USA; Department of Proteomics, The Novo Nordisk Foundation Center for Protein Research, University of Copenhagen, Faculty of Heath Sciences, Copenhagen, Denmark; Center for Biomedical Research, Faculty of Medicine, Universidad de los Andes, Santiago, Chile; Instituto Gulbenkian de Ciência, 2780-156 Oeiras, Portugal; Instituto Medicina de Molecular João Lobo Antunes, Faculdade de Medicina da Universidade de Lisboa, 1649-028 Lisboa, Portugal

## Abstract

Citrullination, the deimination of arginine residues into citrulline, has been implicated in the aetiology of several diseases. In multiple sclerosis (MS), citrullination is thought to be a major driver of pathology, through hypercitrullination and destabilization of myelin. As such, inhibition of citrullination has been suggested as a therapeutic strategy for MS. Here, in contrast, we show citrullination by peptidylarginine deiminase 2 (PADI2) is required for normal oligodendrocyte differentiation, myelination and motor function. We identify several targets for PADI2, including myelin-related proteins and chromatin-associated proteins, implicating PADI2 in epigenetic regulation. Accordingly, we observe that PADI2 inhibition and its knockdown affect chromatin accessibility and prevent the upregulation of oligodendrocyte differentiation genes. Moreover, mice lacking PADI2, display motor dysfunction and decreased number of myelinated axons in the corpus callosum. We conclude that citrullination is required for oligodendrocyte lineage progression and myelination and suggest its targeted activation in the oligodendrocyte lineage might be beneficial in the context of remyelination.

## Introduction

The conversion of a peptidylarginine into a peptidylcitrulline can introduce profound changes in the structure and function of the modified protein. This posttranslational modification, also known as protein citrullination or deimination, is calcium dependent and is mediated by the family of enzymes called peptidylarginine deiminases (PADIs)^1^. Five PADI isozymes (PADI1, 2, 3, 4 and 6) have been identified in mammals, and their expression pattern and function is tissue-specific^2^. Citrullination has been recently shown to play key roles in multiple cellular processes such as inflammatory immune responses^3^, apoptosis^4^, and regulation of pluripotency^5^. In the central nervous system (CNS), PADI2 is the most prevalent expressed PADI and occurs preferentially in oligodendrocytes (OL) and other glial cells^6^. OLs arise at postnatal stages from the differentiation of OL precursor cells (OPCs) and their most prominent function is the formation of the myelin sheath, an electrical insulating layer for the axons, essential for proper neuronal communication and thus CNS function^7^.

PADI2 has been reported to heavily citrullinate myelin binding protein (MBP), a fundamental myelin component. The ratio MBPcit/MBPtotal is high in the first four years of life and similarly in multiple sclerosis (MS), a disease where demyelination occurs^8^. Upon citrullination, MBP arginine residues lack positive charges, thereby having a reduced interaction with the negatively charged lipids. Consequently, citrullinated MBP does not form compact sheaths and the myelin remains immature or becomes unstable^9^. In accordance, transgenic mice overexpressing PADI2 in OL cells display myelin loss^10^. In light of this, PADIs, and in particular PADI2, have been thought to have a detrimental effect in MS and thus an improvement of the disease symptoms can be achieved by inhibiting their actions^11^. Indeed, treatment of a MS mouse model Experimental Autoimmune Encephalomyelitis (EAE) with 2-chloroacetamidine (2-CA), a PADI2 and −4 inhibitor leads to disease attenuation^12^. The improvement on disease progression by 2-CA treatment was not specifically assigned to the lack of PADI activity in OL lineage cells alone but to inhibition of all PADI-expressing cells, including immune cells that play an essential role in the disease development.

Interestingly, however, no changes in disease progression were observed upon EAE induction in PADI2 KO mice^13^. This could point to a possible compensation of PADI2 functions by other PADIs, or to additional functions of PADI2 in EAE disease progression. While citrullination of myelin might play an important role in MS, other proteins such as histone H3 have been found to be citrullinated in MS^14^. Thus, the role of PADI2-mediated citrullination in MS might not be confined to myelin. Furthermore, while the effects of *Padi2* overexpression in OL have been characterized, its absence in OL lineage cells has not been further investigated, nor its physiological function in OL lineage cells and its significance for the myelin integrity maintenance.

## Results

### *Padi2* expression is increased upon OL differentiation

By analyzing our single-cell RNA-Seq dataset of the OL lineage in the adult and juvenile mouse brain^15^, we identified *Padi2* as the predominant *Padi* expressed in OLs (Supplementary Fig. 1a). Interestingly, *Padi2* expression is found in OPCs, increases in committed OL precursors (COPs) and newly formed OLs (NFOLs) and peaks at more mature stages (Supplementary Fig. 1a). Surprisingly, we did not observe expression of *Padi4*, which has been previously suggested to be present in myelin^14,16^. To further investigate the pattern of expression of *Padi2* during OL lineage progression, we cultured OPCs isolated from postnatal day P1-P4 brains of the transgenic mouse line *Pdgfra*-histone 2b (H2B)-GFP^17^ in which GFP expression is under the control of the endogenous *Pdgfra* promoter locus. GFP+ OPCs were collected with fluorescence activated cell sorting (FACS) to plates and expanded in media containing the growth factors (GFs) bFGF and PDGFAA and differentiated into OLs by removing the GFs for 2 days (Fig. 1a). Gene expression of the differentiation markers *Mbp* and *Sox10* was upregulated and the progenitor marker *Pdgfra* was reduced upon GFs removal (Fig. 1b). In agreement with the single-cell RNA-seq data, *Padi2* was expressed in OPCs and it was greatly enhanced upon differentiation (Fig. 1b and Supplementary Fig. 1b). To investigate *Padi2* expression in the OL lineage in vivo, we isolated OPCs and OLs from the brain of the transgenic mouse line *Pdgfra*-Cre-loxP-GFP^18^ in which all OL lineage cells express the GFP reporter (Fig. 1c). GFP+ OPCs were collected with FACS from P1-P4 pups (all GFP+ cells are OPCs since OLs are not formed yet) and GFP+ CD140-OLs were collected with FACS from juvenile (P21) and adult (P60) mice (at this stage GFP+ cells comprise all OL progeny including CD140+ OPCs). As a confirmation of the sorted populations we have assessed the expression of *Pdgfra* and *Mbp* as markers for OPCs and differentiated oligodendrocytes, respectively (Fig. 1d). *Padi2* mRNA was substantially enriched in OLs from both juvenile and adult brains when compared to postnatal OPCs (Fig. 1d). At the protein level, and in agreement with our gene expression data, we observed a continuous increase in PADI2 protein from P1 to adult in the spinal cord of wild type mice, concomitant with the increase in the OL marker MBP (Fig. 1e). Thus, PADI2 is rapidly upregulated upon OPC differentiation, suggesting a role of this citrullinating enzyme at this stage of OL lineage progression.

**Fig. 1.**
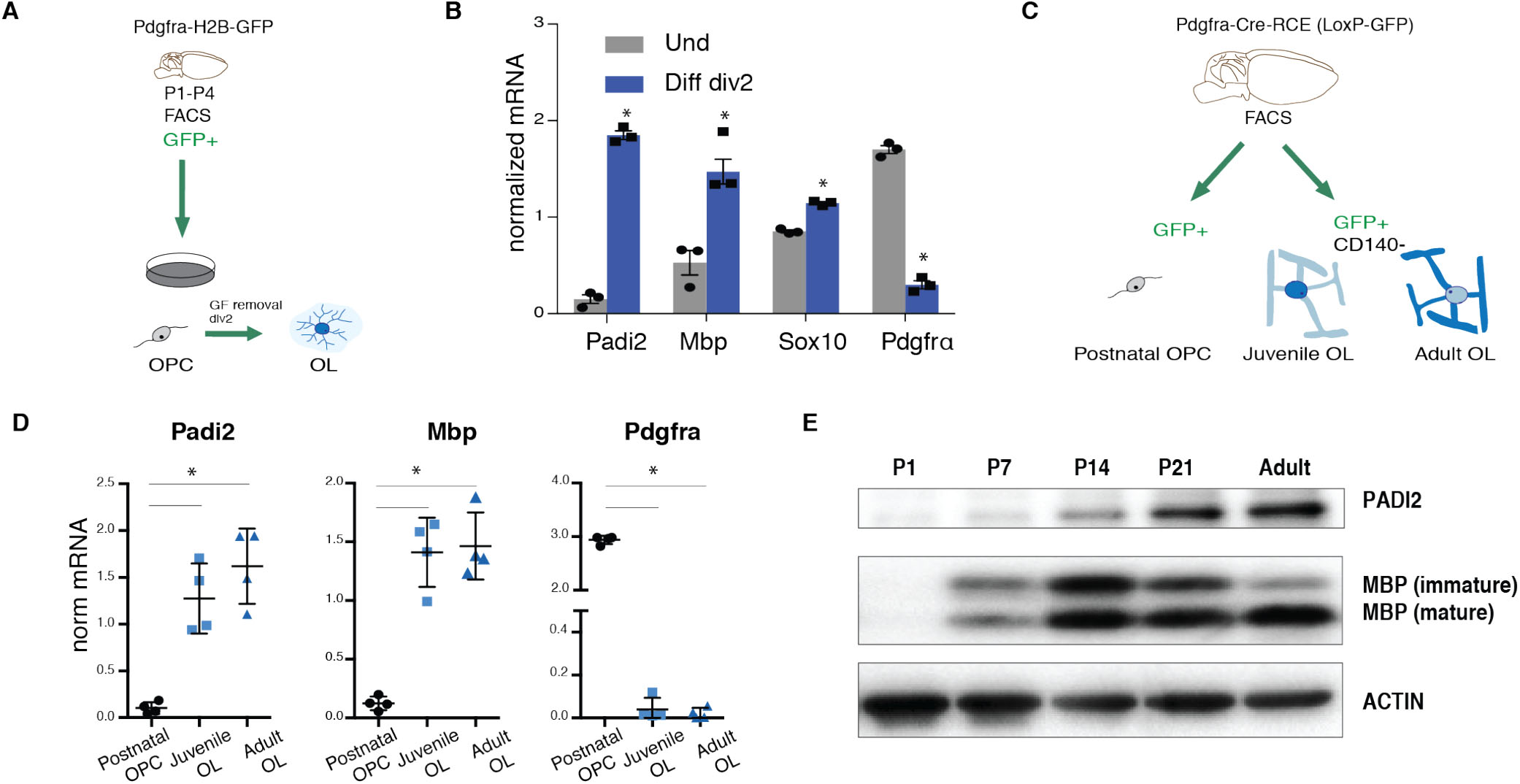
*Padi2* expression is substantially increased upon OL differentiation. (A) Schematic representation of the methodology used for OPC in vitro cultures, P1-P4 GFP+ OPCs are dissociated from brains of the transgenic mouse line *Pdgfra*-H2B-GFP and FACS sorted to plates to expand in the presence of growth factors (GF), GF are removed to induce differentiation for 2 days. (B) Comparative gene expression analysis of OPCs and 2-days differentiated OLs. n=3, two-tailed t-test *p<0.05, error bars represent SEM. (C) Schematic representation of the methodology used to specifically isolate OPCs, juvenile and adult OL from the postnatal (P1-P4), juvenile (P21) and adult (P60) brains of the transgenic mouse line *Pdgfra*-Cre-loxP-GFP; GFP+ cells were depleted of the OPC marker CD140a to specifically isolate OLs. (D) Comparative gene expression analysis of OPCs, juvenile and adult OLs. n=4, one-way ANOVA non-parametric, *p<0.05, error bars represent SEM. (E) Western blot for PADI2 and MBP on the spinal cords of P1, P7, P14, P21 and adult wild type mice, ACTIN signal is an internal loading control.

### Reduction of PADI2 activity hinders OL differentiation

In order to investigate whether the increased expression of PADI2 upon OL differentiation has a functional significance, we inhibited PADI activity in the mouse OPC cell line Oli-neu^19^ and rat primary OPCs, using Cl-amidine, a pan-inhibitor of PADI enzymes. Strikingly, Oli-neu cells failed to differentiate in the presence of this inhibitor, as observed by the lack of cell branching typical of differentiating OLs shown with CNPase immunostaining (Fig. 2a). In accordance, gene expression analysis on undifferentiated and differentiated Oli-neu cells treated with Cl-Amidine revealed a striking impairment on OPC differentiation. The increase of *Mbp*, *Sox10* and *Mog* mRNA levels, crucial for proper OPC differentiation, was no longer observed after Cl-Amidine during differentiation (Fig. 2b). Interestingly, the expression of the transcription factor (TF) *Sox9*, required for OL specification^20^ and downregulated upon OPC differentiation^15^, was increased upon PADI inhibition (Fig. 2b). Similar results were obtained by treating Oli-neu with another PADI-inhibitor 2-CA (Supplementary Fig. 1c). Importantly, these effects were also observed in rat OPC primary cultures, where under differentiating conditions, the treatment with Cl-Amidine impaired differentiation as observed by the lower mRNA expression levels of the differentiation markers *Mog* and *Mag* (Fig. 2c).

**Fig. 2.**
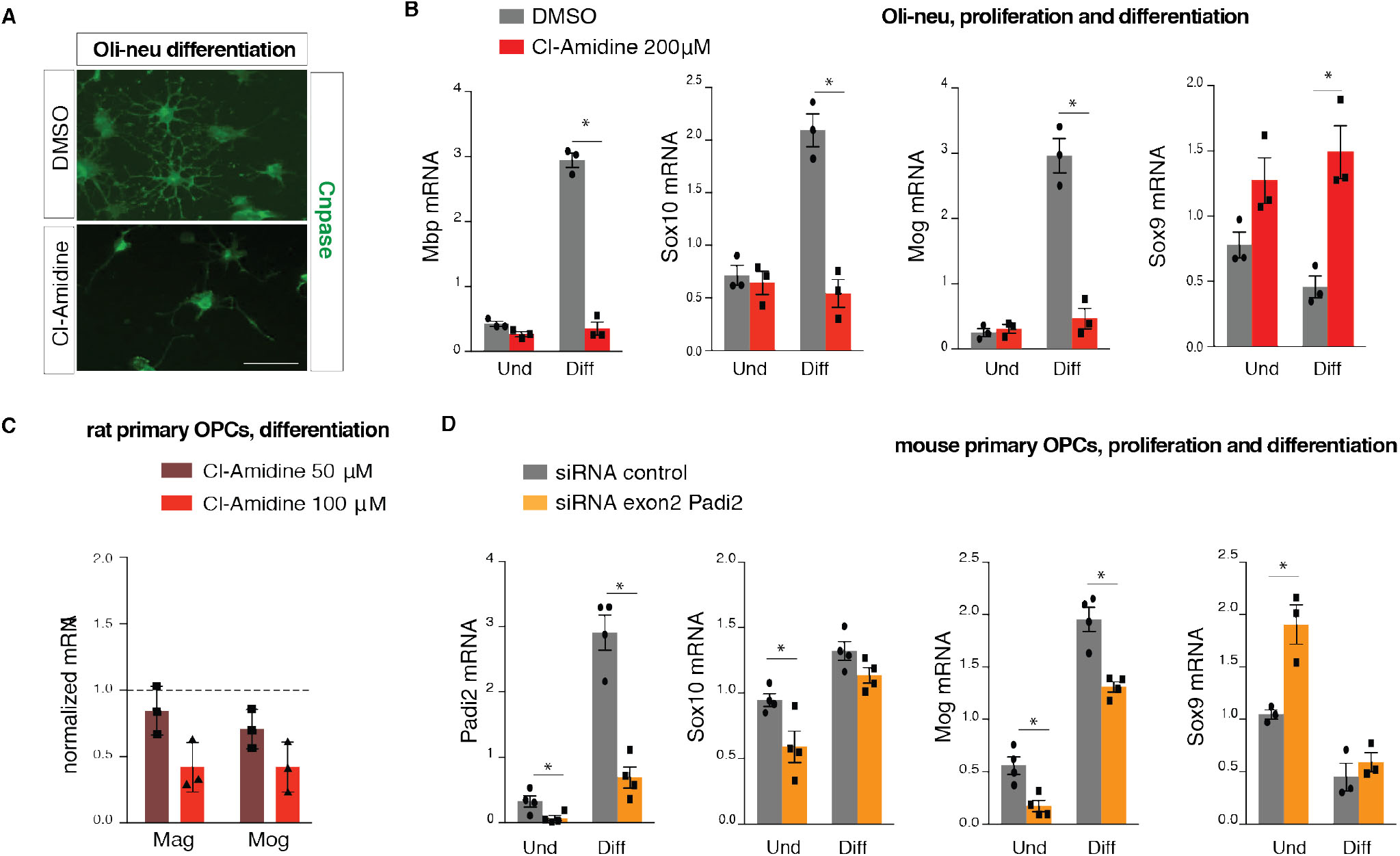
PADIs inhibition and *Padi2* knockdown hinders OPC differentiation. (A) Immunocytochemistry for CNPase on 2-days differentiated Oli-neu upon 2 days treatment with DMSO or Cl-Amidine 200μΜ. (B) Comparative gene expression analysis of proliferating/undifferentiated (Und) Oli-neu cells and 2 days differentiated Oli-neu cells (Diff) treated with DMSO or Cl-Amidine 200μM for 2 days, n=3 two-tailed t-test *p<0.05, error bars represent SEM. (C) Gene expression analysis on proliferating rat OPCs treated with Cl-Amidine 50μM and 100μM, dashed line represents control levels of *Mag* and *Mog*. n=3, Wilcoxon matched pairs test, error bars represent SD. (D) Comparative gene expression analysis on undifferemtiated (Und) and 2-days differentiated OLs (Diff) transfected with either scrambled siRNA or *Padi2* siRNA targeting exon2, n=4 two tailed t-test *p<0.05, error bars represent SEM.

To investigate whether the effect of PADI inhibitors was due to PADI2 inhibition, we specifically decreased *Padi2* mRNA levels in mouse primary FACS OPCs by transfecting cells with siRNA against *Padi2* (targeting exon 2). Similar to PADI inhibition, the reduction of *Padi2* expression led to a decrease in the mRNA levels of *Sox10* and *Mog* and an increase in *Sox9* (Fig. 2d). Mog mRNA levels were also reduced in *Padi2* knockdown conditions after 2-days of differentiation (Fig. 2d). Likewise, *Padi2* knockdown in Oli-neu cells mimicked the effects of PADI inhibition (Supplementary Fig. 1d). Thus, our findings indicate that PADI2 is dramatically increased on the onset of OL differentiation, where it contributes to the transition from an OPC state to a NFOL state.

### OLs are reduced in conditional *Padi2* knockout mouse lines

To investigate if the observed effects of PADI2 have an impact on OL differentiation and myelin maintenance in vivo, we generated a *Padi2* conditional knock-out (cKO) mouse line, *Pdgfra-Cre;* RCE:loxP; *Padi2*−/− where *Padi2* is not present in any cells derived from *Pdgfra*+ cells, which include the whole OL lineage. We estimated the number of CC1+ cells per area at P21, in both corpus callosum and in the dorsal funicullus of the spinal cord. In accordance to our results in Oli-neu and primary FACS OPC cultures (Fig. 2), we observed a reduction in the number of CC1+ OLs in the spinal cord of *Padi2* cKO mice, compared to littermate controls (Fig. 3a, b). Nevertheless, we did not observe such an effect in the corpus callosum (Fig. 3a). This finding led us to further investigated if these OLs exhibited the same levels of OL-related genes by performing qPCR on the brains of P21 controls and *Padi2* cKO. We observed that *Pdgfra*, *Mbp* and *Mog* mRNA were unaltered in FACS cKO GFP+ cells (Supplementary Fig. 2c), indicating that the effect of *Padi2* cKO is more prominent in posterior regions of the CNS.

**Fig. 3.**
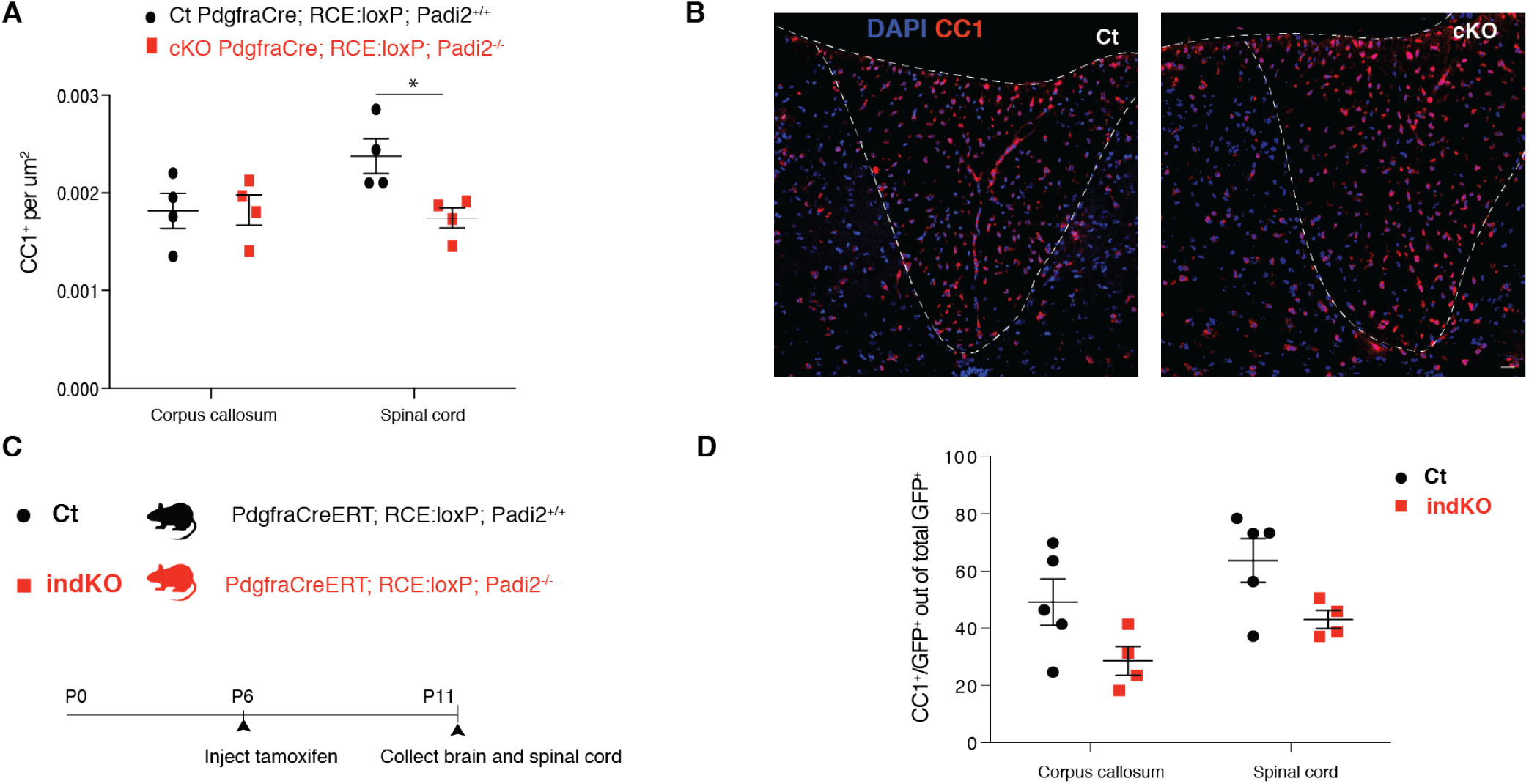
*Padi2* knockout mice displays decreased OPC differentiation. (A) CC1 + cells are presented as number of positive cells per area in corpus callosum and in spinal cord of *PdgfraCre*; RCE-loxP; *Padi2*+/+ and *PdgfraCre*; RCE-loxP; *Padi2*−/− mice. n=4, error bars represent SEM. (B) Representative images of the spinal cord dorsal funicullum of *PdgfraCre;* RCE-loxP; *Padi2*+/+ and *PdgfraCre;* RCE-loxP; *Padi2*−/− mice stained with CC1 antibody and DAPI. scale bar= 20μm. (C) Schematic representation of the strategy used to deplete *Padi2* immediately before the peak of OPC differentiation: *PdgfraCreERT*; RCE-loxP; *Padi2*+/+ and *PdgfraCreERT*; RCE-loxP; *Padti2*−/− mice were injected with tamoxifen at postnatal day 6 (P6) and sacrificed at P11, brains and spinal cords were collected. (D) The number of CC1 and GFP double positive cells were estimated out of total GFP+ cells both in corpus callosum and in spinal cord of *PdgfraCreERT*; RCE-loxP; *Padi2*+/+ and *PdgfraCreERT*; RCE-loxP; *Padti2*−/− mice. n=5 and n=4, two-tailed t-test *p<0.05, error bars represent SEM. Littermate controls were used in all experiments.

We also generated an inducible *Padi2* cKO (indKO) mouse line, *Pdgfra*-CreERT; RCE:loxP; *Padi2*−/−, where *Padi2* could be ablated from OPCs by tamoxifen treatment at early postnatal stages (Fig. 3c). Since we observed that PADI2 mainly modulates the transition between OPCs and NFOLs in vitro (Fig. 2), we depleted *Padi2* specifically at the onset of OPC differentiation by injecting mice with tamoxifen at P6 and collecting the brains and spinal cord at P11 (Fig. 3c). We show a reduction, although not statistically significant (p=0.0828 for corpus callosum and p=0,0567 for spinal cord), of OLs formed upon tamoxifen-induced OPC *Padi2* depletion, as assessed by the percentage of CC1+ cells out of total GFP+ in the corpus callosum and in the spinal cord of *Padi2* indKO animals, compared to littermate controls (Fig. 3d). Thus, our results in two independent mouse lines where *Padi2* is ablated from the OPC stage suggest that PADI2 is important for the generation of OLs in vivo.

### *Padi2* knockout mice display impaired motor and cognitive functions, and a decrease in the number of myelinated axons

Given the decrease in OLs upon *Padi2* cKO in the OL lineage and the targeting of myelin proteins by PADI2, we investigated whether *Padi2* cKO mice presented a behavioral phenotype, in comparison with controls. Interestingly, *Padi2* cKO mice exhibited impairment in motor coordination and balance as assessed by decreased latency to fall in the rotarod test (Fig. 4a). Moreover, we observed similar impairments in the full *Padi2* KO (Padi2 fKO), where exon 1 of *Padi2* is deleted in all cells^13^. In addition, we found an increase in the number of slips on the beam test in the *Padi2* cKO mice, strengthening the motor impairment phenotype (Fig. 4b). Strikingly, when testing the *Padi2* cKO mice for cognitive functions, these mice performed worse in the novel object recognition test, displaying a decrease in the discrimination index that provides an indication of recognition memory (Fig. 4c).

**Fig. 4.**
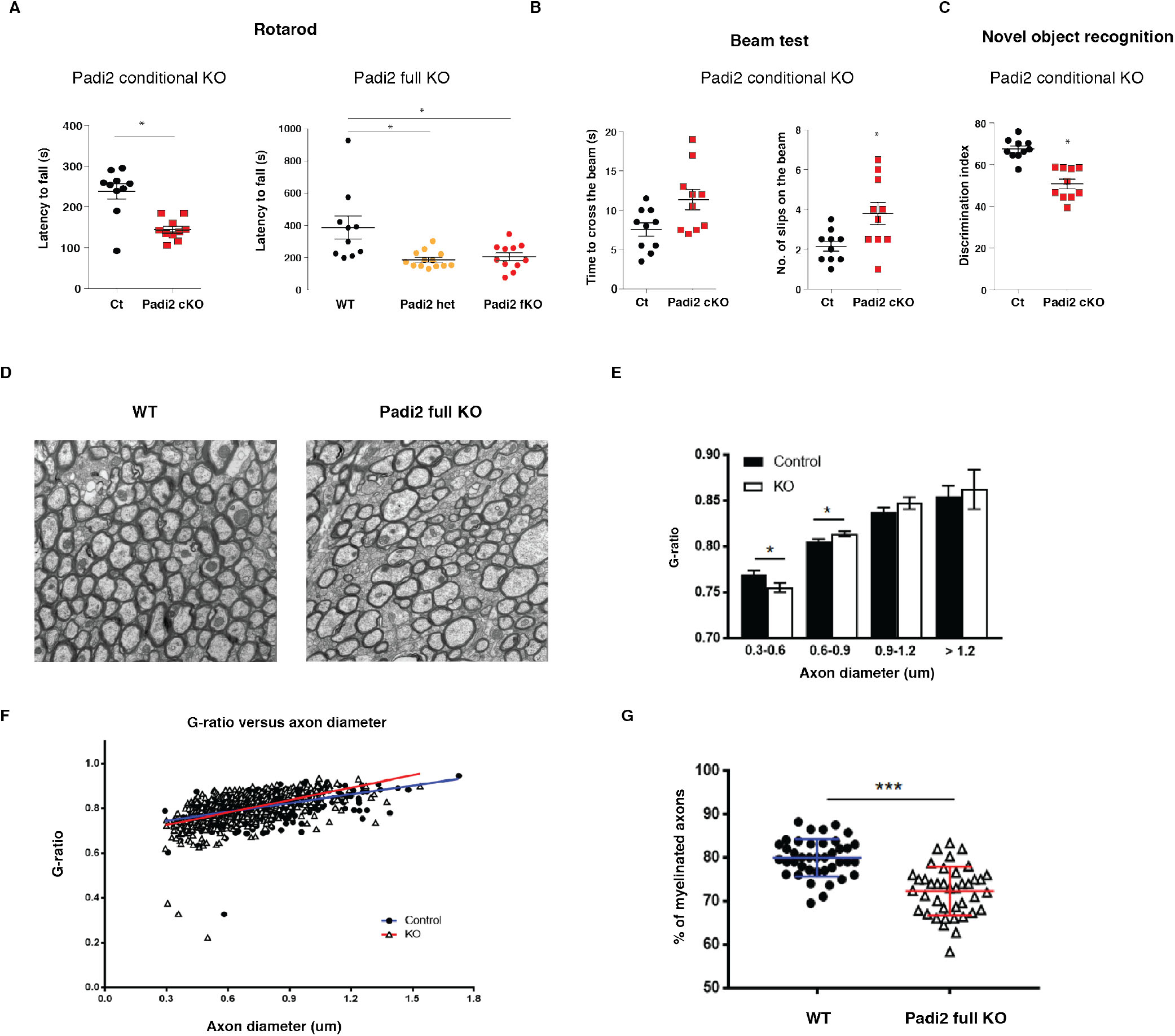
*Padi2* conditional and full knockout mice exhibit impaired motor and cognitive functions and decreased myelinated axons. (A) (B) and (C) Behavior tests for motor (rotarod and beam test) and cognitive (novel object recognition) performance were done in controls (*PdgfraCre*; RCE-loxP; *Padi2*+/+ and and wild type WT), *Padi2* heterozygous (*Padi2* het) and *Padi2* full (fKO) and conditional knockouts (*Padi2* cKO: PdgfraCre; RCE-loxP; *Padi2*−/−). For the rotarod test the time spent on rotating rod, latency to fall, was measured in seconds (s), and for the novel object recognition test the discrimination index corresponds to percentage ratio between the time exploring novel object and the total time spent exploring both objects. Wt n=8, *Padi2* het n=11, *Padi2* fKO n=11, one way ANOVA *p<0.05, error bars represent SEM. Ct n=10 and *Padi2*cKO n=10 Mann-Whitney test *p<0.05, error bars represent SEM. (D) Representative EM images for Wt and *Padi2* fKO corpus callosum. (E) and (F) Scatter plots of g-ratio as a function of axon diameter (μm) and graph plot for the g-ratio distribution across different axon diameters in wt and *Padi2*fKO. (G) Percentage of myelinated axons present in the corpus callous of Wt and *Padi2*fKO. Mann-Whitney U tests were performed for EM analysis.

To further investigate the mechanisms underlying the impact of PADI2 on motor and cognitive functions we have analyzed the myelin structure of the *Padi2* fKO mice and respective controls by Electron Microcopy (EM). We found a significant decrease in the number of myelinated axons in the corpus callosum of adult mice together with a decrease in the g-ratio of small diameter axons and an increase in the g-ratio of axons with 0.6-0.9 μm diameter (Fig. 4d-f). Moreover, the percentage of myelinated axons was decreased in *Padi2* fKO mice compared to controls (Fig. 4g). As such, our results indicate that PADI2 plays an important role for efficient OL differentiation and myelination, as evidenced by the motoric and cognitive defects occurring upon its ablation.

### PADI2 is present in both nucleus and cytoplasm in OL lineage cells

To disclose the possible mechanisms of action underlying the observed in vitro and in vivo effects, we have further investigated the cellular location of *Padi2* in OL lineage cells. Despite that PADI4 is the only one displaying a nuclear localization signal (NLS), PADI2 has recently been detected in the nucleus of mammary epithelial cells^21^ and was described to be able to citrullinate nuclear histone H3R26 in MC7 breast cancer cells^22^. Remarkably, immuno-cytochemistry analysis showed the presence of PADI2 protein both in the nucleus and in the cytoplasm of OPCs (MBP-negative) and OLs (MBP-positive) (Fig. 5a). In order to confirm the localization of PADI2 in the nucleus, we transfected Oli-neu cells with the fusion vector *Padi2*-ZsGreen. We observed the presence of the green fusion protein both in the nucleus and cytoplasm of these cells (Fig. 5b), indicating that PADI2 can indeed be nuclear in OL lineage cells and might thereby regulate transcription through epigenetic mechanisms.

**Fig. 5.**
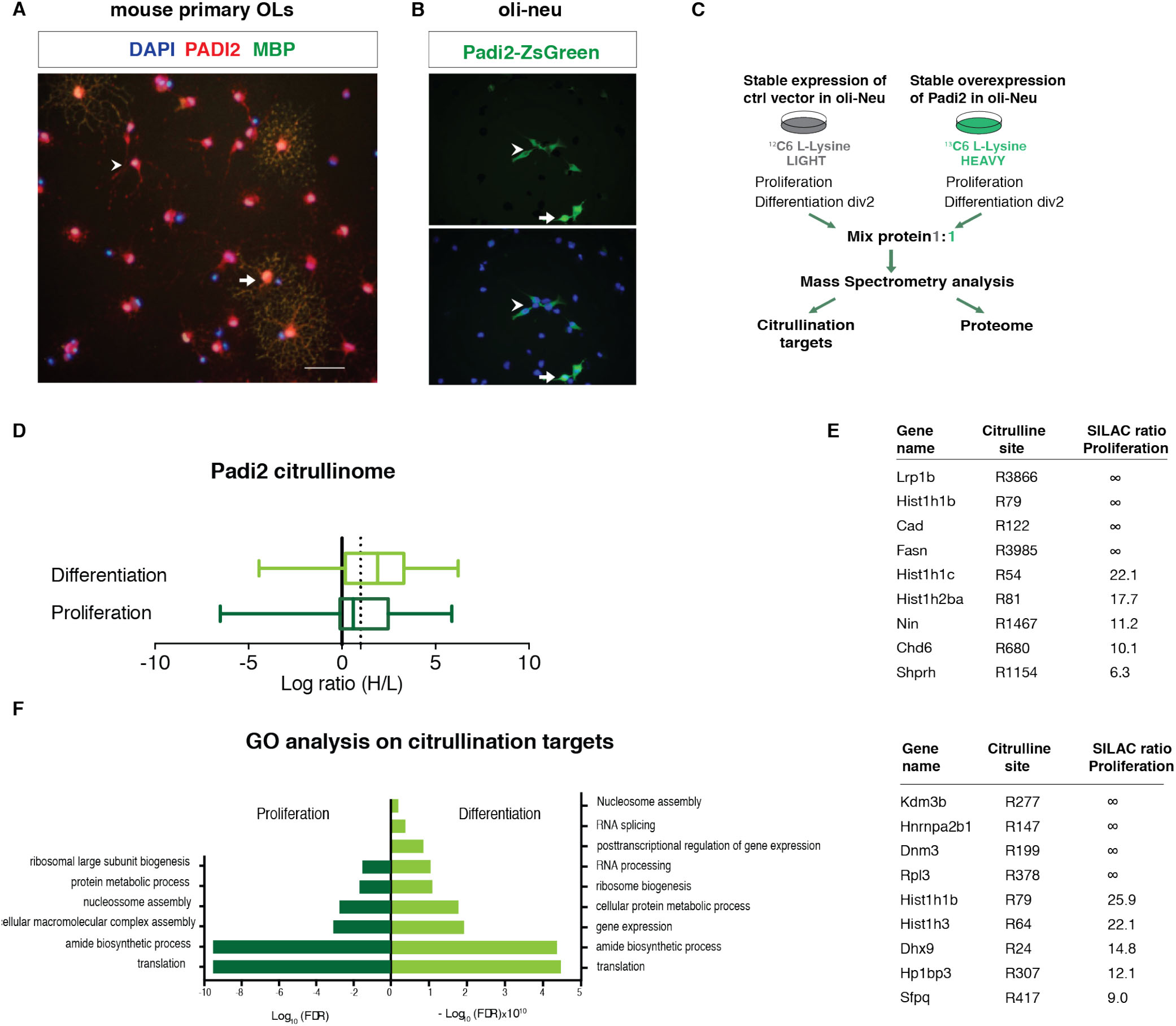
*Padi2* overexpression unveils new citrullination targets. (A) Immunocytochemistry in 2-days differentiated OLs for PADI2 (in red) and MBP (in green) depicting PADI2 in the nucleus and cytoplasm of MBP postitive (arrow) and MBP negative (arrowhead) cells (B) Transfection of Oli-neu cells with *Padi2* fused with ZsGreen shows the PADI2-ZsGreen both in the cytoplasm (arrowhead) and in the nucleus (arrow). (C) Schematic representation of the strategy used to determine the citrullination targets by means of SILAC followed by Mass Spectrometry. Control and overexpressing *Padi2* cell lines are fed with 12C LIGHT and 13C HEAVY media, respectively. Proteins from proliferating and 2-days differentiating cells are collected from both LIGHT and HEAVY media and mixed 1:1 to further mass Spectrometry analysis. (D) Box plot representing the fold change of the identified citrullination targets in Oli-neu cells both in proliferation or differentiation conditions. Ratios are represented as log2 (ratio H/L) and PADI2 mediated citrullination targets were considered above a threshold of 1 (dashed line). (E) Table representing top citrullination targets of PADI2 in proliferation and differentiation conditions. ∞ represent proteins where only peptides with HEAVY labeling were detected.(F) Gene ontology for biological process analysis on the PADI2 citrullinated proteins. The most significant categories (+/- Log10 (false discovery rate)) were plotted.

### PADI2 citrullinates nuclear proteins

To determine what could be the molecular targets of PADI2 in both nuclei and cytoplasm, we generated a stable *Padi2* overexpressing Oli-neu cell line, where citrullination activity was greatly enhanced when tested in an Antibody-based assay for PADI activity (ABAP)^23^ (Supplementary Figure 2a), and performed quantitative proteomics, by applying stable isotope labeling of amino acids in cell culture (SILAC) followed by mass spectrometry^24^. SILAC relies on the incorporation of a light (12C) or a heavy (13C) isotopic version of lysine into proteins of control and overexpressing *Padi2* cell lines, respectively. The proteomes of fully light (L) and heavily (H) labelled cells were mixed 1:1 and the relative dissimilarities in the proteome citrullination target sites of these cell populations was determined and quantified by mass spectrometry (Fig. 5c). Interestingly, we observed a striking difference in the number of citrullination targets and ratios between proliferating and differentiating Oli-neu cells (Fig. 5d). In the proteome of proliferating Oli-neu, we found up to 151 citrullination sites, of which 81 were upregulated (H/L SILAC ratio higher than 1.5) with *Padi2* overexpression. In contrast, up to 500 citrullination sites were identified in differentiating Oli-neu cells, of which 347 were upregulated upon *Padi2* overexpression (Fig. 5d and Supplementary Table 1). These observations indicate that OLs have an intracellular environment that is more permissive to PADI2 activity than OPCs.

We also uncovered citrullination sites arising only with *Padi2* overexpression (no H/L SILAC ratio with peptides detected only in heavy proteomes; Fig. 5e and Supplementary Table 1). Consistent with our previous observations on the localization of PADI2 in the OL lineage (Fig. 4a, b), PADI2 targets were not only cytoplasmic such as ribosomal proteins and myelin proteins (for example HSPA8, CNP and RAB1, ENO1 and EEF1A25), but also nuclear (Fig. 5e and Supplementary Table 1). Chromatin components and modulators such as histone H1 (HIST1H1B), histone H2b (HIST1H2BA), histone H3 (HIST1H3), the histone demethylase KDM3B and the chromodomain helicase DNA binding protein 6 (CHD6), among others, were identified to be citrullinated by PADI2 at specific arginines (Fig. 5e). We also observed citrullination of several RNA binding proteins, consistent with the abundance or arginines in their RNA binding motifs (Fig. 5e). Gene ontology (GO) analysis indicated that proteins citrullinated by PADI2 are involved in translation, ribosomal biogenesis and nucleosome assembly both in proliferating and differentiating conditions, and RNA splicing, RNA processing and posttranscriptional regulation of gene expression in differentiating conditions (Fig. 5f and Supplementary Table 4). Thus, PADI2 activity is not circumscribed to myelin and cytoplasmic proteins but is also relevant to nuclear processes.

### PADI2 protein-protein interaction network is enriched in cytoplasmic and myelin proteins

Our data suggests that PADI2 might be involved in different biological cell processes occurring both in the nucleus and in the cytoplasm. To further explore the PADI2 role in OL lineage cells, we identified the PADI2 protein interactors by pulling-down biotin-tagged PADI2. Control (expressing biotin ligase, BirA, and empty vector) and biotin-tagged *Padi2* (bio*Padi2*) Oli-neu cell lines (expressing BirA and the biotin-tagged *Padi2*) were fed with 12C or 13C isotopes, respectively, immunoprecipitated with streptavidin beads and combined for further mass spectrometry analysis (Fig. 6a). As expected, PADI2 was the protein enriched the most (Fig. 6b). We uncovered up to 74 shared PADI2 protein interactors in proliferating and 2-days differentiated Oli-neu cells (Fig. 6b, c, e and Supplementary Table 2), among which nuclear proteins such as elongator protein complex 1 (ELP1). Nevertheless, the number of interactors was much lower than citrullination targets, suggesting that most of the PADI2 interactions leading to citrullination are transient and cannot be captured. Yet, GO analysis on the biological processes showed common categories with the GO analysis of the PADI2 citrullination targets, such as translation and posttranscriptional regulation of gene expression (Fig. 6d), suggesting once again a role for PADI2 in these processes. Interestingly, one of the categories in the GO analysis, the intracellular protein transport, comprises RAN and KPNB1, two key regulators for active nuclear transport, which suggests that these proteins can be involved in the PADI2 nuclear-cytoplasmic shuttling (Supplementary Table 2). When performing GO analysis for the cellular component, we also have identified a myelin sheath component of PADI2 interactors (Fig. 6e, myelin sheath partners highlighted in red, Supplementary Table 4). Furthermore, comparison with a previously published myelin proteome^25^ uncovers additional myelin proteins that interact with PADI2 such as WDR1, FASN, among others (Fig. 6e and Supplementary Table 2). Thus, while interactions of PADI2 with nuclear proteins might be more transient, we observed several myelin proteins interacting with this enzyme.

**Fig. 6.**
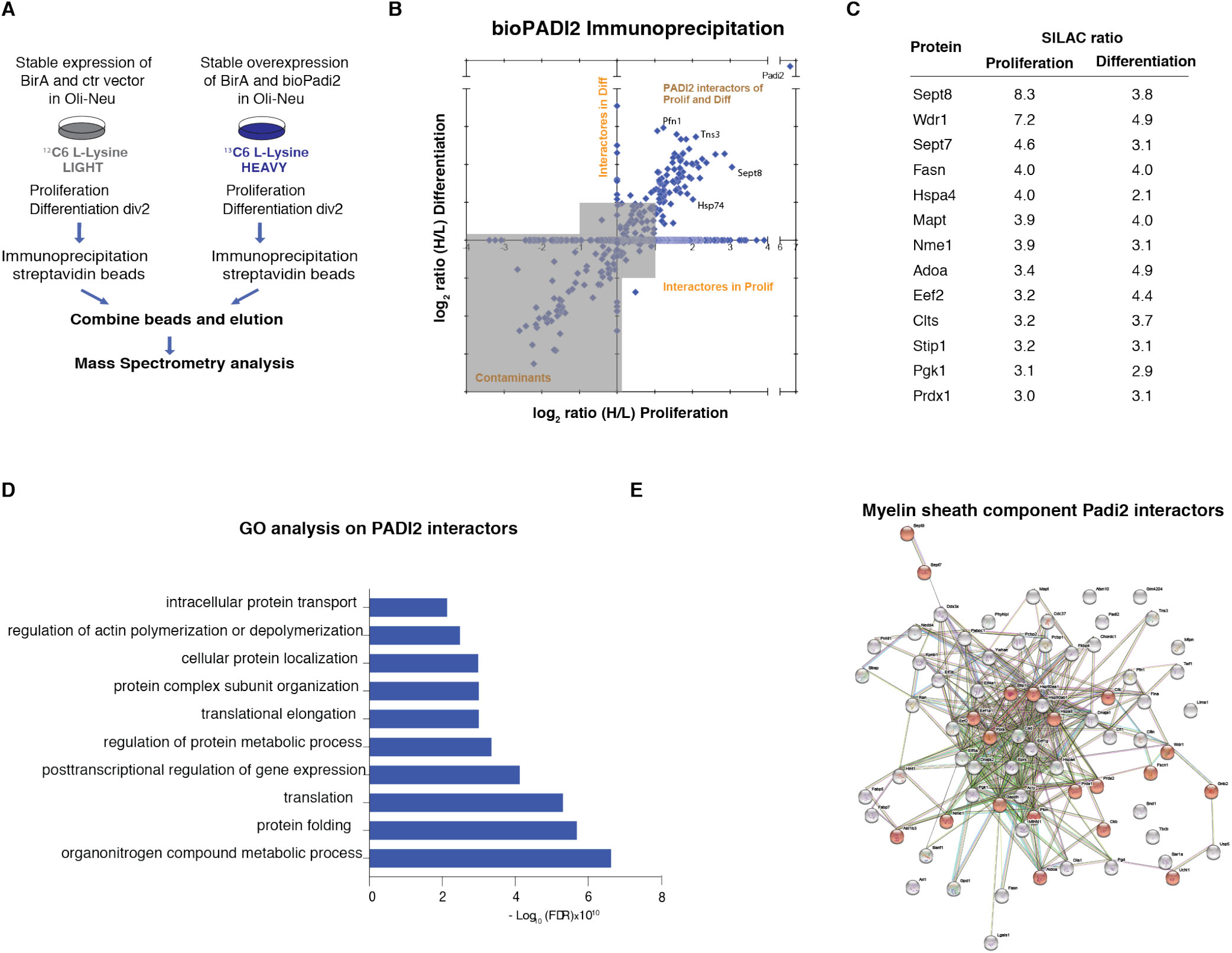
PADI2 protein interactors in Oli-neu cells. (A) Schematic representation of the strategy used to uncover PADI2 interacting proteins by means of SILAC followed by Mass Spectrometry. Control cells (expressing biotin ligase BirA and empty vector) and biotin-tagged *Padi2* (bio*Padi2*) cell lines (expressing BirA and the biotin tagged *Padi2*) are fed with 12C LIGHT or 13C HEAVY media, respectively. Immunoprecipitation (IP) with streptavidin beads is performed on the same amount of proteins collected from control or bio*Padi2* cells. After IP the streptavidin beads from the two conditions are mixed are analyzed by Mass Spectrometry. (B) Graph plot representing all detected immunprecipitated proteins by Mass Spectrometry in both proliferating and upon 2-days differentiation of Oli-neu cells. Ratios are represented as log2 (ratio H/L) and PADI2 interacting proteins were considered for analysis when displaying a threshold above 1 and detected in both proliferation and differentiation (all targets in the up right quadrant). (C) Short list of the 10 top PADI2 interacting proteins that were previously shown in the myelin proteome. (D) Gene ontology for biological process analysis on the PADI2 interacting proteins. The most significant categories (+/- Log10 (false discovery rate)) were plotted. (E) Network analysis of all PADI2 interactors, proteins found on the cellular component GO analysis for the myelin sheath are highlighted in red.

### Histone citrullination by PADI2 is associated with regulation of OL differentiation genes

Since we found PADI2 in the nucleus of OL lineage cells and identified several chromatin-associated proteins as targets of PADI2, we investigated whether citrullinated nuclear proteins, such as histones, are present in OL lineage cells, using an orthogonal technique. Indeed, western blot analysis of histone citrullination shows the presence of histone H3R26Cit and histone H3R2+8+17Cit in proliferating Oli-neu cells (Fig. 7a). Treatment with 200 μM of the pan PADI inhibitors Cl-Amidine or 60 μM of 2-CA led to a reduction on these modifications (Fig. 7a). We also observed an increase of histone H3R2+8+17Cit and Histone H3R26Cit in the *Padi2* overexpressing cell lines indicating that PADI2 is mediating histone H3 citrullination and thus acts as an epigenetic regulator in OL lineage cells (Fig. 7b). While the expression of *Mog* and *Sox9* was not altered upon *Padi2* overexpression, we observed upregulation of *Mbp* and *Cnp* in differentiating cells (Fig. 7c), corroborating their downregulation upon PADI2 inhibition/*Padi2* knockdown (Fig. 2). Thus, histone citrullination might be one of the mechanisms by which PADI2 regulates the transition of OPCs to NFOLs.

**Fig. 7.**
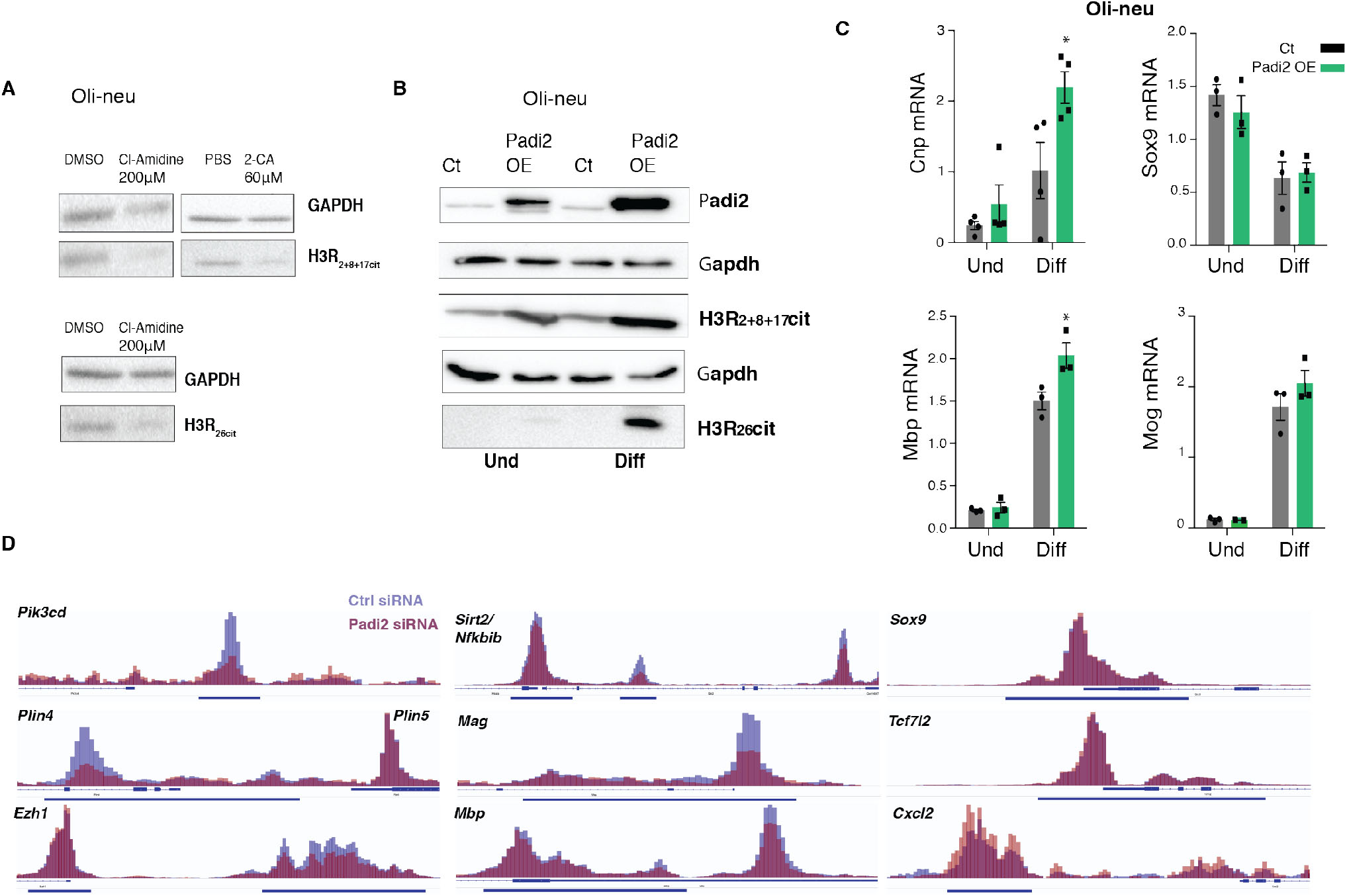
PADI2 citrullinates histones and regulates the expression and chromatin acessibility of genes involved in OL differentiation. (A) Western blot for H3R2+8+17cit and H3R26cit upon treatment of proliferating Oli-neu cells with either DMSO or Cl-Amidine 200μM for 2 days, GAPDH signal is an internal loading control. Images represent n=2. (B) Western blot analysis for PADI2, Histone H3R2+8+17cit and H3R26 on *Padi2* overexpressing cell line versus control. GAPDH was used as a loading control. Images represent n=3. (C) Comparative gene expression analysis of proliferating/undifferentiated (Und) Oli-neu cells and 2 days differentiated Oli-neu cells (Diff) overexpressing *Padi2*, n=3 (*Mbp, Mog, Sox9*) n=4 (*Cnp*) two-tailed t-test *p<0.05, error bars represent SEM. (D) IGV genome browser overlay views depicting chromatin accessibility (assessed by ATAC-Seq) near transcription start sites, in Oli-neu transfected with Ctrl siRNAs (blue) and siRNA against *Padi2*(red) (n=3, samples pooled for visualization, same scale for ctrl siRNA and *Padi2* siRNA). Regulatory regions (promoters/enhancers marked with H3K27ac or H3K4me3 in Oli-neu or OPCs, see Methods) are highlighted with blue horizontal bars.

### *Padi2* knockdown results in reduced accessibility at OL differentiation genes

To further examine the possible role of PADI2 in transcriptional regulation and the chromatin structure, we investigated if the knockdown of *Padi2* would affect the chromatin accessibility. We performed assay for transposase-accessible chromatin using sequencing (ATAC-seq)^26^ to identify changes in accessibility of regulatory regions in differentiating Oli-neu upon knockdown of *Padi2* with siRNAs (Fig. 7d). Interestingly, we observed a general reduction in accessibility upon knock down of *Padi2* (Fig. 7d, Supplementary Fig. 2d and Supplementary Table 3). We further investigated accessibility of OL enhancers and promoters, presenting H3K27ac in Oli-neu cells (see methods) or H3K27ac/H3K4me3 in OPCs^27^. Indeed, we observed decreased chromatin accessibility in regulatory regions of genes as *Pik3cd*, *Plin4* or *Ezh1*. Importantly, we also observed such reduction in accessibility in important regulators of OL myelination, including *Sirt2*, *Mag* and *Mbp* (Fig. 7d), and in *Plp1*, *Sox10* and *Mog*, although to minor extent (Supplementary Fig. 2d). Accordingly, gene ontology analysis indicated an enrichment in regulatory regions of genes involved in lipid metabolism (Supplementary Fig. 2e and Supplementary Table 4). Nevertheless, PADI2 appears to target specific regulatory regions, since we did not find regulation of chromatin accessibility at regulatory regions of other genes modulating OL differentiation, such as *Sox9*, *Tcf7l2*, (Fig. 7d), *Id1* or *Olig2* (Supplementary Fig. 2d). Interestingly, we also find a cohort of genes where accessibility of their regulatory regions was instead increased by *Padi2* knockdown, including *Cxcl2* (Fig. 7d), *Hspb8* or *Gabra6*. Gene ontology analysis indicated involvement in processes chemotaxis, Polycomb activity and apoptosis, among others (Supplementary Fig. 2d and Supplementary Table 4). As such, recruitment of PADI2 to specific regulatory regions in OLs regulates their accessibility, which will also ultimately modulate transcriptional output, as observed for *Mog* and *Sox10*.

## Discussion

The formation of myelin is a fundamental biological process that encompasses subsequent steps of OPC specification, differentiation and myelin sheath formation/wrapping of axons. Our study highlights PADI2 as an important player in these last two steps by 1) facilitating OPC differentiation and influencing the OL epigenetic landscape through histone citrullination and 2) interacting with and mediating the citrullination of several myelin protein components. While *Padi2* expression in OLs and its occurrence in the myelin were described before^16^, we report for the first time its presence in OPCs and the dramatic upregulation of *Padi2* in the transition from OPCs to NFOLs and its function facilitating OL lineage progression. To uncover PADI2 mechanisms of action, we further determined the citrullination targets and the binding partners of PADI2. Our data revealed new PADI2 citrullinated proteins and interactors, both located in the nuclear and cytoplasmic compartments. We also demonstrate, using 3 different *Padi2* knockout mouse lines, that PADI2 is important for the OL development and motoric and cognitive behaviour. Our findings indicate that PADI2 mediated citrullination is required for efficient OL development with acting not only as a myelin modulator, through MBP citrullination, but also displaying other functions in the cytoplasm and nucleus, for instance as an epigenetic modulator. Additional mechanisms underlying PADI2 actions in OPC differentiation remain to be investigated.

By using an unbiased proteomics methodology based on SILAC followed by mass spectrometry we have revealed hundreds of novel citrullination protein targets and the respective arginine locations, some of those in core histones such as HIST1H1, HIST1H2 and HIST1H3. PADI2 mediated H3R26Cit has been shown to lead to local chromatin de-condensation and targeted transcriptional activation in breast cancer cells^28^. In a similar manner, when we knockdown *Padi2* we observe a general decrease in chromatin accessibility as revealed by ATAC-seq. Furthermore, upon PADIs inhibition or *Padi2* knockdown we observed a decrease in the pro-differentiation and myelin genes such as *Mbp*, *Mog* and *Sox10* suggesting that PADI2 might be locally enhancing transcription through histone H3 citrullination.

Interestingly, we detected HIST1H1C citrullination on R54 in our cells. This modification was previously described to induce global chromatin decondensation in iPS cells^5^ and to be specifically mediated by PADI4, which suggests that there is an overlap of the substrates of PADI2 and PADI4. Moreover, H3R2+8+17Cit was also induced upon *hPadi4* over-expression^5^. hPADI2 and hPADI4 were shown to exhibit different substrate preferences and hPADI4 had more pronounced substrate specificity, in both Cos-1 and HEK cells or by adding recombinant hPADIs to cell lysates^29^. Additional hPADI2 targets were disclosed by overexpressing hPADI2 in HEK cells using a different technology^30^. Although these are targets for hPADI2 in different human cell types, we observed many common targets with our mouse OL lineage cells such as ribosomal proteins and splicing factors. Thus, PADI2 might have a widespread and undetermined role in basic cellular processes such as mRNA splicing and translation.

Differentiated Oli-neu cells displayed considerably more citrullination targets, higher citrullination activity in basal conditions with the ABAP kit, and a more pronounced increase in H3R2+8+17Cit and H3R26Cit upon *Padi2* overexpression suggesting that PADI2 activity can be enhanced upon differentiation. In agreement, most of PADI2 effects are more noticeable upon OPC differentiation. Of notice, PADI activity can also be context dependent, as these enzymes need a high concentration of calcium to become active, and calcium signaling has been shown to be fundamental for OPC migration, differentiation and myelination^31^. Although several citrullination targets found in proliferating and 2-days differentiated Oli-neu were common and/or showed additional citrullinated arginine in differentiation, they did not overlap completely, indicating that there is a preference for the targeted arginine between these conditions that could be due for instance, to differences in the binding partners. Likewise, not all proteins found interacting with PADI2 were citrullinated and vice versa. Nevertheless, GO analysis for the biological processes on the PADI2 interactors also reveals categories similar to the citrullination targets, such as translation and posttranscriptional regulation of gene expression reinforcing once again a putative role for PADI2 in these processes.

Mice overexpressing *Padi2* under the *Mbp* promoter display abnormal myelination with structural changes of the myelin^10^. This effect was attributed to the increased citrullination of MBP and also of histones that would ultimately cause OL apoptosis^10^. However, a striking finding in our study is the abundance of myelin component proteins within the interactors of PADI2, that might also mediate the effect of PADI2 on myelin when overexpressed^10^. Citrullination of myelin proteins is naturally occurring, and thus normal citrullination levels must certainly have a physiological function. 2’,3’-Cyclic-nucleotide 3’-phosphodiesterase (CNP), which has an important role in the generation of cytoplasmic channels in OLs^32^, displayed PADI2-mediated citrullination of several arginines (Table S1). It is possible that hypocitrullination of CNP upon knocking out PADI2 could lead to disruption of cytoplasmic channels and ultimately to the observed defects on myelination in our *Padi2* fKO mice (Fig. 7). Further investigation would be necessary to determine which citrullination targets of PADI2 in the myelin could be responsible for these defects.

We found that PADI2 absence in OL lineage cells caused decreased OL cell density in spinal cord and a decrease in the number of myelinated axons in the corpus callosum of adult mice, that were reflected in motor and cognitive dysfunctions. Since *Pdgfra* and *Padi2* can also be co-expressed in other non-CNS cells such as Schwann cells, the impairment on motor functions could also be due to the absence of PADI2 in peripheral cells. However, overexpression of *Padi2* did not have any effect on the peripheral myelin^10^ and the EM and OL densities analysis in the brain and spinal cord (Fig. 7) indicate a phenotype in OLs. In addition, the cognitive behavior test also supports a CNS specific PADI2 effect. Analysis of single-cell RNA-Seq data (Supplementary Fig. 1) suggested that PADI2 is the only member of the PADI family robustly expressed in the OL lineage. Although PADI4 was identified by IHC in the myelin^16^ we could not detect *Padi4* mRNA in any OL lineage cell tested nor it was identified in the myelin proteome^25^. However, we did detect very low mRNA levels of *Padi3* and *Padi1* in Oli-neu cells and found it expressed in very few cells in OPCs/OLs collected from P90 brains (data not shown). As such, we cannot totally rule out that these PADIs have a function in OL lineage progression and that compensatory mechanisms might be operational upon PADI2 knockdown, knockout or inhibition, particularly since PADIs can have overlapping substrates.

PADI2 has been long investigated in the context of MS since MBP hypercitrullination is a hallmark of the disease^33^. Our study demonstrates that PADI2-mediated citrullination is also required for OL lineage progression and myelination and uncovers several new citrullination targets in the OL lineage. Thus, targeted activation of PADI2 in the OL lineage might be beneficial in the context of remyelination in diseases as multiple sclerosis.

## ACKNOWLEDGEMENTS

We would like to thank Tony Jimenez-Beristain, Alessandra Nanni, Ahmad Moshref and Johnny Söderlund, and the Clinical Proteomics Mass Spectrometry at Karolinska University Hospital and Science for Life Laboratory for providing assistance in mass spectrometry and data analysis, and Science for Life Laboratory, the National Genomics Infrastructure (NGI), and Uppmax for providing assistance in massive parallel sequencing and computational infrastructure. We thank Belinda Panagel and the CMB FACS facility for all the support in FACS sorting of all cells, and Jean-Jacques Medard and Roman Chrast for help with electron microscopy. We thank Helena Francis, Iris Muller, Marjan Abbasi, Theresa Mader, Diana Hernandez and Wouter Beenker, all students that had participated and pushed forward this project. We thank P. Soriano (Mount Sinai, New York) for the Pdgfra-H2B-GFP mouse. The bioinformatics computations were performed on resources provided by the Swedish National Infrastructure for Computing (SNIC) at UPPMAX, Uppsala University. A.M.F. was supported by the European Committee for Treatment and Research of Multiple Sclerosis (ECTRIMS). Work in P.C.’s research group was funded by NIH-NINDS to PC R01-NS052738. Work in G.C.-B.’s research group was supported by Swedish Research Council, European Union (FP7/Marie Curie Integration Grant EPIOPC, Horizon 2020 Research and Innovation Programme/ European Research Council Consolidator Grant EPIScOPE, grant agreement number 681893), Swedish Brain Foundation, Ming Wai Lau Centre for Reparative Medicine, Swedish Cancer Society (Cancerfonden), Petrus och Augusta Hedlunds Foundation and Karolinska Institutet. Raw data has been deposited in GEO GSE115929. This manuscript was formatted using the BioRxiv template in the Overleaf software (Author: Ricardo Henriques, License: Creative Commons CC BY 4.0).

## AUTHOR CONTRIBUTIONS

AMF and GCB designed all experiments. AMF, MM, and JL performed the in vitro experiments, AMF and AH performed the tissue collection and microscopic analysis, PR and AS performed the behavior experiments and PE and PC oversaw, AS, MF and RK processed and analyzed Electron Microcopy data and PC oversaw, MV-G was involved in all the cloning, SCL and MLN performed and analyzed Mass Spectrometry data. EA analyzed the ATAC-Seq experiments, while AR analyzed the Chip-Seq experiments, under supervision of DC. GCB and AMF wrote the paper, with contribution from all authors, and were involved in all the data analysis.

## Experimental procedures

### Animals

All mice were maintained under a pathogen-free environment at the animal facility. All experimental procedures were performed following the guidelines and recommendations of local animal protection legislation and were approved by the local committee for ethical experiments on laboratory animals (Stockholms Norra Djurförsöksetiska nämnd in Sweden). The following transgenic mice strains were used: *Pdgfra*-H2BGFP knock-in mice^17^ where H2BeGFP fusion gene is expressed under the promoter of *Pdgfra*; *Pdgfra*-Cre BAC transgenic mice (The Jackson Laboratories, CA, USA), where Cre recombinase is under the control of the *Pdgfra* promoter; RCE:loxP-GFP reporter mice (Gord Fishell, NYU Neuroscience Institute); *Pdgfra*-CreERT BAC transgenic mice (The Jackson Laboratories, CA, USA), *Padi2*tm1a(KOMP)Wtsi (UCDAVIS KOMP Repository Knockout Mouse Project) and *Actb*:FLPe (The Jackson Laboratories, CA, USA). All individual mice strains were in a C57BL/6 genetic background, at the exception of RCE:loxP-GFP that was in a CD1 background. *Padi2*fl/fl was generated by breeding *Padi2*tm1a(KOMP)Wtsi with Flp deleter mice. *Padi2*fl/fl mice were crossed with RCE:loxP-GFP, *Pdgfra-*Cre or *Pdgfra*-CreERT. Tamoxifen (Sigma T5648) was dissolved in corn oil and administrated i.p. in P6 *Pdgfra-*CreERT;RCE:loxP;*Padi2*fl/fl pups at a dose of 2mg/40g. Brain and spinal cord were subsequently collected at P11.

Use of full *Padi2* KO mice was strictly compliant with the guidelines set forth by the US Public Health Service in their policy on Human Care and Use of Laboratory Animals, and in the Guide for the Care and Use of Laboratory Animals. Mice were maintained under a pathogen-free environment at the animal facility of Mount Sinai Medical Center. All procedures received prior approval from the Institutional Animal Care and Use Committee (IACUC). The C57BL/6-*Padi2-*KO mice were obtained from the Scripps Research Institute (Kerri Mowen’s laboratory) which was originally from Lexicon Genetics (Woodlands, TX).

### Behavioral analysis

For the beam balance test the fine motor coordination and balance was assessed by beam walking assay. Performance on the beam was quantified by measuring the time taken by the mouse to traverse the beam and the number of paw slips that occurred in the process. The beam apparatus consisted of a PVC plastic beam (1.2 cm wide × 100 cm long) located 50 cm above the table top on two poles. A black box was placed at the end of the beam as the finish point. Nesting material and faeces from home cages was placed in the black box to attract the mouse to the finish point. A nylon or cotton pad was placed below the beam, to cushion any falls. Mice were trained three times a day for two consecutive days until they were able to cross the beam without hesitation. No data were collected during these sessions. During the test session, the time needed to cross the beam and the number of paw slips were recorded. The beam and box were cleaned of mouse droppings and wiped with 70% ethanol before the next animal was placed on the apparatus. The novel object recognition task was conducted as described previously^34^, and was used to assess the non-spatial hippocampal memory. A mouse was presented with two similar objects during the first session, and then one of the two objects was replaced by a new object during a second session. The amount of time taken to explore the new object provided the index of recognition memory. The task consisted of habituation, training, and testing phases conducted on separate days. During habituation, mice were allowed to freely explore an empty white box (60 cm wide × 60 cm long × 47 cm high) for 5 min/day for two days. Twenty-four hours later, mice were re-habituated (familiarization session) in the same box for 10 min, with two identical objects placed inside, 5 cm away from the walls. Mice were allowed to freely investigate until they accumulated a total of 20 sec exploring the objects. Exploration and recognition of the object were determined when the nose of the animal was in contact with an object or directing nose to the object within a defined distance (<2 cm). Mice were then immediately returned to their home cage. After the familiarization session, the objects and the open field was cleaned with 70% ethanol to minimize olfactory cues. After 1 hour of familiarization session, object recognition was tested, using the same procedure as in training except that one of the familiar objects was substituted with a novel object (cleaned and free of olfactory cues). The position of the novel object (left or right) has to be randomized between each mouse and each group tested. Time spent with each object was recorded using a stopwatch by a trained observer. Mice inherently prefer to explore novel objects; thus, a preference for the novel object indicates intact memory for the familiar object. A discrimination index was calculated according to the following formula: time of novel object exploration/time of novel plus familiar object exploration×100. Rotarod behavioral analysis was performed for evaluation of coordination and motor learning. Mice were trained on a Rotarod (Ugo Basile, Comerio VA, Italy) at a constant speed (4 rpm), 120 s per day for 3 consecutive days. Subsequently, test day consisted of three trials on an accelerating rotarod for up to 300 s (starting at 4 rpm accelerating to a final speed of 40 rpm), with mice returned to their home cages for at least 30 min between trials. The time at which each animal fell from the rod was recorded as a measure of motor coordination and repeated measurements of speed at the time of animal falling were averaged.

### Electron Microscopy

For EM analysis, deeply anesthetized C57BL/6-*Padi2*-KO mice (n=4 per condition) were perfused transcardially using a varistaltic pump (Manostat) with 0.1M PBS (pH 7.4) followed by cold Karnovsky’s fixative (2.0% PFA/2.5% Glutaraldehyde in 0.1M Sodium Phosphate buffer). Using a McIlwain tissue chopper, coronal brain sections between −2.2 and −2.5 bregma were obtained and corpus callosum tissue was micro-dissected and post-fixed overnight in 2.0% PFA/ 2.5% Glutaraldehyde in 0.1M Sodium Phosphate buffer. The samples were then processed using standard protocols employed by the Stony Brook University Microscopy Core (see below, and^35,36^). The following day (after overnight post-fixation), an additional fixation step was carried out using 2% osmium tetroxide in 0.1M PBS (pH 7.4), after which samples were dehydrated in a graded series of ethyl alcohol. Finally, samples were embedded using Durcupan resin in between two pieces of ACLAR sheets (Electron Microscopy Sciences). Ultrathin sections of 80nm were obtained using a Lecia EM UC7 ultramicrotome and placed on formvar coated slot copper grids (Electron Microscopy Sciences). Uranyl acetate and lead citrate counterstained samples were viewed under a FEI Tecnai12 BioTwinG2 electron microscope. Digital images (minimum 10 per animal, taken at 11,000x) were acquired using an AMT XR-60 CCD Digital Camera system and compiled using Abobe Photoshop software. G-ratio was determined by dividing the mean axon diameter without myelin by the same axon diameter with myelin. A minimum of 100 healthy axons were considered per animal. To determine the size distribution of myelinated fibers in corpus callosum, diameters of all measured axons were separated into four ranges (0.3-0.6, 0.6-0.9, 0.9-1.2, & >1.2 µm). No axons with diameters smaller than 0.3 µm were considered. Data are expressed as comparison of means between the control and knock-out for each range. Percentage of myelinated axons was determined by placing a centered grid over the image using ImageJ software, and counting whether the axon at each intersection was myelinated or not. A minimum of 100 axons was counted per image, and 10 images were quantified per animal. All measurements were acquired on electron microscopy sections images using an ImageJ software. Mann-Whitney U tests were performed to assess statistical differences between control and knock out mice for both differences in g-ratio per axon diameter range and percent myelinated axons.

### Tissue dissociation

Cells were isolated from postnatal, juvenile and adult brains from *Pdgfra*-Cre;RCE:loxP mice, and from juvenile brains from *Pdgfra*-Cre;RCE:loxP;*Padi2*+/+ and *PdgfraCre*;RCE:loxP;*Padi2*fl/fl mice with Neural (for postnatal) and Adult (for juvenile and adult) Tissue Dissociation kits (P) (Miltenyi Biotec). Cell suspensions were then filtered with 30µm filter (Partec) and FACS sorted for selection of GFP+. FACS sorted cells were collected in eppendorf tubes, centrifuged at 400xg, resuspended in Qiazol and frozen for further RNA extraction/cDNA synthesis.

### FACS

GFP+cells were FACS sorted using a BD FACSAria III Cell Sorter B5/R3/V3 system. Cells were collected either in PBS for further RNA extraction processing or in OPC media directly into coated plates. For OPC removal on OL lineage cells from juvenile and adult brains, CD140 (anti-mouse CD140 APC conjugated, BD Bioscience) labelling was performed in tissue-isolated cells and GFP+CD140+ cells were excluded. To define the gates for specific CD140+ cells we have performed an additional labelling with APC conjugated IgG (BD Bioscience) as a negative control.

### Cell culture

Primary OPC culture: GFP+ OPCs were obtained with FACS from P1-P4 brains from the *Pdgfra-*H2BGFP mouse line. The mouse brains were removed and dissociated in single cell suspensions using the Neural Tissue Dissociation Kit (P) (Miltenyi Biotec, 130-092-628) according to the manufacturer’s protocol. Cells were seeded in polyL-lysine (O/N) (Sigma P4707) plus fibronectin (1h) (Sigma F1141) coated dishes and grown in proliferation media comprising DMEM/Gmax (life tec 10565018), N2 media (life tec 17502048), Pen/Strep (life tec 15140122), NeuroBrew (Miltenyi 130-093-566), bFGF 20ng/ml (Peprotech 100-18B) and PDGF-AA 10ng/ml (R&D 520-BB-050). For OPC differentiation, cells were left for 2 days in medium without bFGF and PDGF-AA. Oli-neu cell culture: Oli-neu cells were grown in poly-L-lysine coated dishes and expanded in proliferation media consisting of DMEM (life tec 41965062), N2 supplement, Pen/Strep Glu (life tec 10378016), T3 (sigma T6397) 340ng/ml, L-thyroxine (sigma 89430) 400 ng/ml, bFGF 10ng/ml and Pdgf-BB 1ng/ml. For Oli-neu differentiation, media without growth factors was added for two days in vitro and in the presence of 1µM of erb inhibitor (PD174265, St Cruz, sc-204170). Rat OPC cultures: primary rat OPCs were isolated as reported previously^37^. Briefly, mixed glia cultures were first derived from rat brain cortex at postnatal day 1. OPCs were selected by the A2B5 antibody followed by purification with the magnetic rat anti-mouse IgM beads (Miltenyi Biotec). Purified OPCs were cultured in the SATO medium supplemented with PDGFA and bFGF, and were induced to differentiate with T3 replacing the growth factors. PADI inhibition: Olineu cells were treated either with Cl-Amidine (Millipore 506282) or 2-chloroacetamidine (2CA) (Sigma C0267) at a concentration of 200 µM and 60 µM for one and two days, respectively, in proliferation and differentiation media.

### SILAC

To generate LIGHT and HEAVY labeled cell lines, Oli-neu cells were grown and expanded in LIGHT (Lysine 12C) or HEAVY (Lysine 13C) media for at least five passages. The media was prepared with SILAC Advanced DMEM/F12-FLEX kit (Life Tec, MS10033) and contained 0.1% of dialyzed FBS, 10ng/ml bFGF, 10ng/ml PDGF-BB, 400ng/ml of L-thyroxine and 340ng/ml of T3. For the differentiation, growth factors and the FBS were removed. LIGHT labeled cells lines correspond to all controls, scramble over-expression control and BirA control, while HEAVY labeled cell lines comprise the overexpression of *Padi2* or the biotin tagged *Padi2*. A control experiment in which control cells were also fed with heavy media and then mixed 1:1 with light control cells was performed in order to confirm that all proteins in heavy conditions were heavily labeled (Supplementary Fig. 2b).

### *Padi2* overexpression cell lines and knock down

Padi2 and Ctr overexpression cell lines were generated by insertion of the following vectors with piggyBac transposon system: pB-CAG-Ctr, pB-CAG-*Padi2* and for the control and overexpression of *Padi2*, respectively; pB-CAG-BirA together with pB-CAG-IRES-GFP as a control and pB-CAG-BirA together with pB-CAG-bio*Padi2*-IRES-GFP for the biotin tagged *Padi2*. The gateway system was used to clone all these vectors (Gateway^®^ LR Clonase^®^ II Enzyme mix, 11791020 and Gateway^®^ BP Clonase^®^ II Enzyme mix11789020) in Oli-neu cells. pB-CAG-Ctr, pBCAG-Padi2, pB-CAG-BirA + pB-CAG-IRES-GFP and pBCAG-BirA + pB-CAG-bio*Padi2*-IRES-GFP were transfected together with piggyBac transposase (pBase) expression vector by lipofection according to the manufacturer’s instructions (Lipofectamine^®^ 2000, Invitrogen 11668019). To select for biotin tagged *Padi2* and pB-CAG-IRES-GFP vectors cells were FACsorted for GFP, all the other cell lines express the hygromycin resistance gene in the DNA integrated vectors and were selected and expanded in media containing 200 µg/ml of hygromycin. For the expression of the fusion *Padi2* plasmid in Oli-neu OPCs, *Padi2* was cloned into the pZsGreen1-N1 plasmid (Clontech, 632448). Oli-neu and primary mouse OPCs were then transfected with these plasmids with Lipofectamine2000 according to the manufacturer’s instructions. For silencing *Padi2* we used the siDESIGN center from Dharmacon. The following siRNAs were selected and purchased from Dharmacon GE Healthcare: mouse *Padi2*; sense: 5’ CGUACGUGAUGGAGAGGCAUU 3’; antisense: 5’ UGCCUCUCCAUCACGUACGUU 3’ and ONTARGETplus Non-targeting siRNA #1 (D-001810-01-20). Knockdown experiments for *Padi2* were performed both in Oli-neu and mouse primary OPCs with Lipofectamine2000, following manufacturer’s instructions. For mouse primary OPCs, upon 4h of adding the lipids:DNA complexes to the cells, media was changed to either proliferation or differentiation and cells were collected two days after. For Oli-neu experiments, upon 4h of adding the lipids:DNA complexes to the cells, media was changed to proliferation for one day and cells were either collected at this point (proliferation condition) or changed to differentiation media one more day (differentiation condition).

### ABAP

Cells were collected in lysis buffer [20mM Tris-HCl, pH 7.4; 100mM NaCl; 10mM β-mercaptoethanol; 10% glycerol; protease inhibitor complete (Roche)] and sonicated for 5 minutes at high power with 30 sec on/off cycles at 4°C. Then the samples were centrifuged at 13000 × g at 4°C for 30 minutes and the supernatant was immediately used for further application. PADI activity was determined with the Antibody Based Assay for PAD activity (ABAP) kit (Modiquest Research, MQ17.101-96) and performed according to manufacturer’s protocol. HRP-conjugated secondary antibody was visualized with a TMB substrate solution [1 mg/ml TMB (Sigma Aldrich); 0.1M sodium acetate, pH 5.2; 0.01% hydrogen peroxide] and the reaction was stopped with sulphuric acid [2M H2SO4]. The absorbance at 450 nm was determined with a FLUOstar Omega plate reader (BMG Labtech). Human PADI4 enzyme was used as control enzyme activity and was diluted in deimination buffer [40 mM TrisHCl, pH 7.5; 5 mM CaCl2; 1mM DTT] with concentrations between 0.002 mU (minimum deimination) and 2.0 mU (maximum deimination) to create a standard curve to correlate activity to the optical density measured at 450 nm.

### RNA extraction, cDNA synthesis and quantitative real-time PCR (qRT-PCR)

RNA was extracted with the miRNeasy mini kit (Qiagen, 217004) for the Oli-neu cells or with the miRNeasy micro kit (Qiagen, 217084) for the primary OPCs and direct tissue isolated cells according to manufacturer’s protocols. Contaminating DNA was degraded by treatment of the samples with RNase-free DNase (Qiagen, 79254) in column. 0.35-1µg of RNA from each sample was reversed transcribed for 1h with the High-Capacity cDNA Reverse Transcription Kit (Applied Biosystems, 4368813) including RNase inhibitor (Applied Biosystems, N8080119). An RT- control was included for each sample. Both the cDNA and the RT- were diluted 1:5 in RNase/DNAse free water. qPCR reactions were run on a StepOnePlus™ System (Applied Biosystems) in duplicate and with reverse transcriptase negative reactions to control for genomic DNA. Fast SYBR^®^ Green Master Mix (Applied Biosystems, 4385616) was used according to the manufacturer’s instructions, each PCR reaction had a final volume of 10µl and 1–2.5µl of diluted cDNA and RT-. The running conditions were 20 seconds at 95°C, followed by 40 cycles of 3 seconds of 95°C and 30 seconds of 60°C, then 15 seconds at 95°C, 1 minute at 60°C and 15 seconds at 95°C. A melting curve was obtained for each PCR product after each run, to control for primer dimers and gene-specific peaks. Random PCR products were also run in a 2–3% agarose gel to verify the size of the amplicon. *Tbp* and *Gapdh* were run as housekeeping genes. Relative standard curves were generated for each gene to determine relative expression (CT values are converted to arbitrary quantities of initial template per sample). Expression levels were then obtained by dividing the quantity by the value of the geometric mean of the housekeeping genes. PCR primer sequences used are listed in table 1.

**Table 1.**
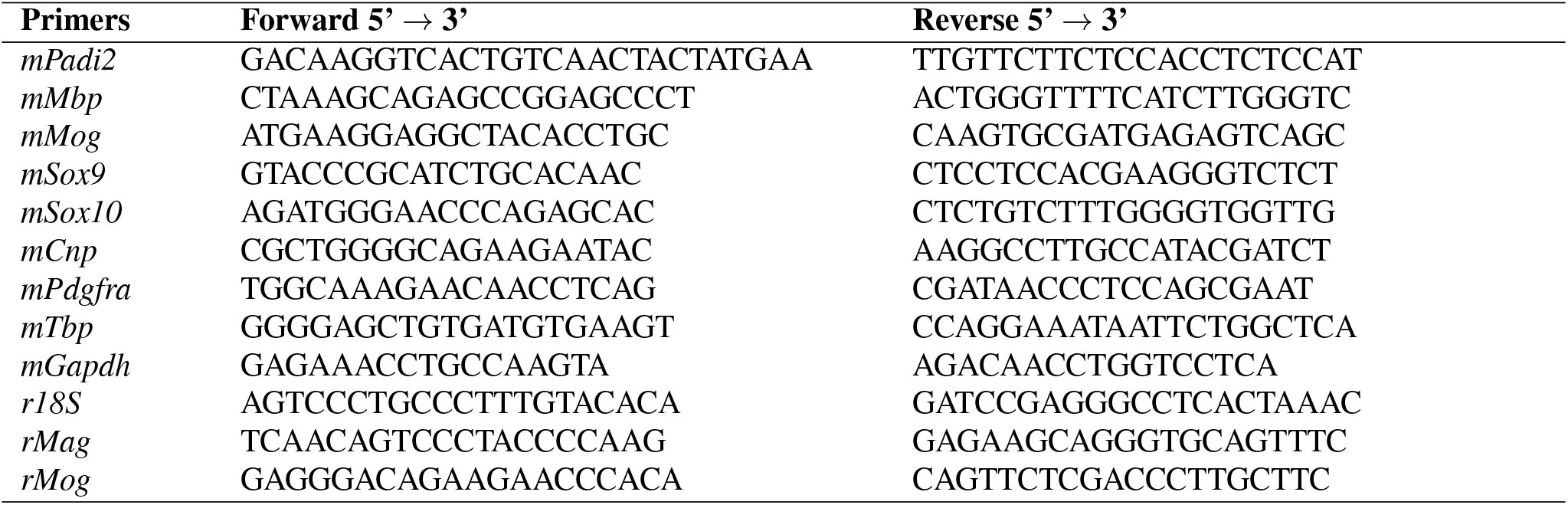
Primers used in qRT-PCR.

### Immunofluorescence

For immunocytochemistry cells were fixed in 4% formaldehyde for 10 minutes, washed in PBS and incubated overnight at 4°C with the primary antibodies anti-CNP (abcam ab6319, 1:200) anti-MBP (Abcam, ab7349, 1:200) in PBS/0.5%Trition/10% normal donkey serum (Sigma, D9663). Cells were washed with PBS and then incubated for 2 hours with Alexa Fluor-conjugated antibodies (Invitrogen, Alexa Fluor anti-mouse 488 1:1000 and Alexa Fluor anti-mouse 555 1:1000). For immunohistochemistry mice were perfused at P11 (for tamoxifen-induced *Pdgfra*-CreERT;RCE:loxP/*Padi2*+/+ and *Pdgfra*-CreERT;RCE:loxP;*Padi2*-/-) or at P21 (for *Pdgfra-*Cre;RCE:loxP;*Padi2*+/+ and *Pdgfra*-Cre;RCE:loxP;*Padi2-*/-) with PBS followed by 4% PFA. Brains and spinal cords were dissected and post-fixed with 4% PFA for 1h, at 4°C. The tissues were then cryo-protected with a 30% sucrose solution for 24 hours. The tissues were embedded into OCT (Tissue-Tek) and sectioned coronally (20 um thickness). Sections were incubated overnight at 4°C in the following primary antibodies: GFP (Abcam, ab13970, chicken 1:2000) and CC1 (anti-APC; Millipore, OP80, Mouse 1:100) diluted in PBS/0.5% Triton/10% normal donkey serum. After washing the section with PBS, secondary Alexa Fluor-conjugated antibodies diluted in PBS/0.5% Triton/10% normal donkey serum were incubated for 2h at room temperature (Invitrogen, Alexa Fluor anti-chicken 488 1:1000 and Alexa Fluor anti-mouse/goat 555 1:1000). Slides were mounted with mounting medium containing DAPI (Vector, H-1200) and kept at 4°C until further microscopic analysis.

### Microscopy and quantitative analysis

To estimate the number of OL (CC1+ cells), newly formed/ Cre recombined OL (GFP+CC1+) and Cre recombined OPCs (GFP+PDGFRa+), 3 sections per animal (at same positions of the anterior-posterior axis) were analyzed. For each section, images were taken for the entire corpus callosum and dorsal funiculus of the spinal cord using a Zeiss LSM700 Confocal. The total number of CC1+ cells and GFP+CC1+ or GFP+PDGFRa+ double positive cells was counted using Image J software. The percentage of OL and OPCs out of total recombined cells at P11 corpus callosum and spinal cord was calculated by estimating the percentage ratio of GFP+CC1+ out of total GFP+cells for OLs and GFP+PDGFRa+ out of total GFP+cells for OPCs. For the estimation of OL density in juvenile mice CC1+cells were divided by the correspondent areas of corpus callosum and dorsal funiculus.

### Western Blot

Cells were collected in 2x Laemmli buffer [120mM Tris-HCl, pH 6.8; 4% SDS; 20% glycerol] and sonicated for 5 minutes at high power with 30 sec on/off cycles at 4°C to shear genomic DNA. Protein concentrations were determined on nanodrop and concentrations were equalized with 2x Laemmli buffer. Bromophenol blue (0.1%) and β-mercaptoethanol (10%) were added to the samples and the samples were boiled at 95°C for 5 minutes to denature the protein. Equal volumes were loaded in a SDS-PAGE for protein separation and transferred (100V for 90’) to a PVDF membrane (GE-healthcare) activated in methanol. The membranes were blocked in blocking solution [TBS; 0.1% Tween 20; 5% milk] for 1 hour at room temperature and incubated overnight with primary antibody (diluted in blocking solution) at 4°C, washed 3 times 10 minutes in TBS-T [TBS;0.1% Tween 20] and incubated with a horseradish peroxidase (HRP)-conjugated secondary antibody (diluted in blocking solution) for 2 hours at room temperature. Proteins were exposed with either enhanced chemiluminescence solution (GE healthcare) or Supersignal west femto (Thermo Scientific) at a ChemiDoc™ XRS imaging system (Bio-Rad). Primary antibodies were used against PADI2 (rabbit polyclonal; 12110-1-AP; ProteinTech; 1:300), GAPDH (rabbit monoclonal; 5174S; Cell Signaling; 1:1000), H3 (mouse monoclonal; ab10799; Abcam; 1:1000) H3Cit2+8+17 (rabbit polyclonal; ab5103; Abcam; 1:1000), H3Cit26 (rabbit polyclonal; ab19847; Abcam; 1:1000). Secondary antibodies were used at a dilution of 1:5000, anti-rabbit (A6667; Sigma), anti-mouse (A4416; Sigma).

### PADI2 immunoprecipitation

Proliferating and 2-days differentiated BirA controls and biotin tagged Padi2 cells were fixed fo 8min at RT in 1% formaldehyde solution by adding 1/10 volume to cell media of freshly prepared 11% formaldehyde solution (formaldehyde 11%, 0.1M NaCl, 1mM EDTA pH8, 0.5mM EGTA pH8 and 50mM Hepes pH8). Glycine was added (final 0.125M) to quench formaldehyde, rinsed twice in cold PBS and scrapped for subsequent centrifugation at 4000g for 10min at 4°C. At this point cell pellets can be stored at −80°C. Cell pellets were then resuspended in 1mL of lysis buffer (10mM Hepes-KOH pH7.5, 100mM NaCl, 1mM EDTA, 0.5mM EGTA, 0.1% Na-Deoxycolate,0.5% Na-lauroylsarcosinate, protease inhibitors) and sonicated. Cell suspension was centrifuged at 14000g for 10min at 4°C to remove the debris. For the immunoprecipitation, protein concentrations were determined in every sample and equal amounts of protein were used for further streptavidin pull-down. 50µL of streptavidin beads (DB Myone Streptavidin T1, Invitrogen) per 5mg of protein were added to the protein suspensions and incubated overnight at 4°C. Magnetic stand was used to precipitate the beads. At this step beads from BirA controls and biotin-tagged Padi2 were combined for the following washes: 4 times in wash buffer 1 (50mM Hepes pH7.6, 1mM EDTA, 0.1% Na-Deoxycolate), 2 times in wash buffer 2 (50mM Hepes pH7.6) and 2 times in wash buffer 3 (20 mM ammonium bicarbonate NH4HCO3). The beads can be stored at −80°C in this step.

### Mass Spectrometry for citrullination targets

#### SDS-PAGE gel

Extracted proteins were resuspended in Laemmli Sample Buffer, and resolved on a 4-20% SDS-PAGE. The gel was stained with Coomassie blue, cut into 20 slices and processed for mass spectrometric analysis using standard in gel procedure^38^. Briefly, cysteines were reduced with dithiothreitol (DTT), alkylated using chloroacetamide (CAA) (PMID: 18511913), and finally the proteins were digested overnight with endoproteinase Lys-C and loaded onto C18 StageTips prior to mass spectrometric analysis.

#### Mass spectrometric analysis

All MS experiments were performed on a nanoscale EASY-nLC 1000 UHPLC system (Thermo Fisher Scientific) connected to an Orbitrap QExactive Plus equipped with a nanoelectrospray source (Thermo Fisher Scientific). Each peptide fraction was eluted off the StageTip, auto-sampled and separated on a 15 cm analytical column (75 µm inner diameter) in-house packed with 1.9-µm C18 beads (Reprosil Pur-AQ, Dr. Maisch) using a 75 min gradient ranging from 5% to 40% acetonitrile in 0.5% formic acid at a flow rate of 250 nl/min. The effluent from the HPLC was directly electrosprayed into the mass spectrometer. The Q Exactive plus mass spectrometer was operated in data-dependent acquisition mode and all samples were analyzed using previously described ‘sensitive’ acquisition method^39^. Back-bone fragmentation of eluting peptide species were obtained using higher-energy collisional dissociation (HCD) which ensured high-mass accuracy on both precursor and fragment ions^40^.

#### Identification of peptides and proteins by MaxQuant

The data analysis was performed with the MaxQuant software suite (version 1.3.0.5) as described^41^ supported by Andromeda (www.maxquant.org) as the database search engine for peptide identifications^42^. We followed the step-by-step protocol of the MaxQuant software suite^43^ to generate MS/MS peak lists that were filtered to contain at most six peaks per 100 Da interval and searched by Andromeda against a concatenated target/decoy^44^ (forward and reversed) version of the IPI human database. Protein sequences of common contaminants such as human keratins and proteases used were added to the database. The initial mass tolerance in MS mode was set to 7 ppm and MS/MS mass tolerance was set to 20 ppm. Cysteine carbamidomethylation was searched as a fixed modification, whereas protein N-acetylation, oxidized methionine, deamidation of Asparagine and Glutamine, and citrullination of Arginines were searched as variable modifications. A maximum of two mis-cleavages was allowed while we required strict LysC specificity. To minimize false identifications, all top-scoring peptide assignments made by Mascot were filtered based on previous knowledge of individual peptide mass error. Peptide assignments were statistically evaluated in a Bayesian model on the basis of sequence length and Andromeda score. We only accepted peptides and proteins with a false discovery rate of less than 1%, estimated on the basis of the number of accepted reverse hits.

### Mass Spectrometry for PADI2 interactome

#### LC-MS/MS sample preparation

The beads were thawed and resuspended in 100µl 25 mM ammonium bicarbonate. The samples were reduced for 1 hour at room temperature by addition of 1 mM DTT, followed by alkylation for 10 minutes in the dark with 5 mM iodoacetamide. The reaction was quenched by the addition of 5 mM DTT. 0.2 µg of Trypsin (sequencing grade modified, Pierce) was added to the samples and digestion was carried out over night at 37°C. After digestion, samples were spun briefly and the supernatant carefully transferred to a glass vial and dried in a SpeedVac and resuspended in 3% acetonitrile, 0.1% formic acid. On third of the original sample was injected for LC-MS/MS analysis.

#### LC-ESI-MS/MS Q-Exactive

Online LC-MS was performed using a Dionex UltiMate™ 3000 RSLCnano System coupled to a Q-Exactive mass spectrometer (Thermo Scientific). Samples were trapped on a C18 guard desalting column (Acclaim PepMap 100, 75um × 2 cm, nanoViper, C18, 5 µm, 100 Å), and separated on a 50 cm long C18 column (Easy spray PepMap RSLC, C18, 2 µm, 100Å, 75 µmx15cm). The nano-capillary solvent A was 95% water, 5%DMSO, 0.1% formic acid; and solvent B was 5% water,5% DMSO, 95% acetonitrile, 0.1% formic acid. At a constant flow of 0.25 µl min1, the curved gradient went from 2%B up to 40%B in 95 min, followed by a steep increase to 100%B in 5 min. FTMS master scans with 70,000 resolution (and mass range 300-1700 m/z) were followed by data-dependent MS/MS (35 000 resolution) on the top 5 ions using higher energy collision dissociation (HCD) at 30-40% normalized collision energy. Precursors were isolated with a 2m/z window. Automatic gain control (AGC) targets were 1e6 for MS1 and 1e5 for MS2. Maximum injection times were 100ms for MS1 and 150-200ms for MS2. The entire duty cycle lasted 2.5s. Dynamic exclusion was used with 60s duration. Precursors with unassigned charge state or charge state 1 were excluded. An underfill ratio of 1% was used.

#### Peptide and protein identification

The MS raw files were searched using Sequest-Target Decoy PSM Validator under the software platform Proteome Discoverer 1.4 (Thermo Scientific) against a mouse database (Uniprot, downloaded on 140320) and filtered to a 1% FDR cut off. We used a precursor ion mass tolerance of 10 ppm, and a product ion mass tolerance of 0.02 Da for HCD-FTMS. The algorithm considered tryptic peptides with maximum 2 missed cleavage; carbamidomethylation (C) as a fixed modification; oxidation (M), and heavy SILAC labelled lysines (Label 13C(6) / +6.020 Da (K)) as dynamic modifications.

#### Gene Ontology analysis

Gene ontology (GO) enrichment analysis was performed using STRING protein-protein interaction networks^45^, and results adjusted for multiple testing using the Benjamini–Hochberg procedure (FDR).

### ChIP-seq

Oli-neu cells were rinsed with cold PBS and cross-linked with 1% formaldehyde (Sigma) for 10 min at room temperature (RT) on a rotating platform. Fixation was quenched with 0.125M glycine for 5 min at RT. Cells were rinsed with cold PBS containing 1x Protease Inhibitors (PI, Sigma, 11873580001) and collected spinning at 1500rpm for 5 min at 4°C. Pellets were resuspended in swelling buffer (PIPES 5mM, KCl 85 mM, Igepal-CA630 1%, PI 1x), incubated 15min on ice followed by a centrifugation of 1300g for 5min. This step was repeated twice and nuclear pellets were resuspended in lysis buffer (1% SDS, 10 mM EDTA, 50 mM Tris-HCl (pH 8.0) and PI 1x) and incubated for 10 min on ice followed by chromatin sonication using a BioruptorTM UCD200 (Diagenode), high frequency, 30 sec ON/30 sec OFF, for 10 min. Protein A/G coated beads (Invitrogen) were washed and resuspended in ChIP dilution buffer (0.01% SDS, 1% Triton, 1.2mM EDTA, 16.7mM TrisHCl pH8, 167mM NaCl and PI 1x). 20µg of chromatin per ChIP was diluted in ChIP dilution buffer and precleared by incubating with 20µl of pre-washed protein A/G coated beads for 1.5h at 4°C on a rotator. Beads were removed and 2µg of the antibody H3K27ac (Millipore, 17-683) was added to the chromatin and incubated overnight at 4°C on a rotator. Incubate 20µl of beads in 1mg/ml of BSA in PBS overnight at 4°C on a rotator. Wash beads in ChIP buffer and add to the chromatin/ab for 1h at 4°C on a rotator. Chromatin/ab/beads were washed 3 times low salt wash buffer (0.1% SDS, 2mM EDTA, 20mM TrisHCl pH8, 1% TritonX-100, 150mM NaCl), once with high salt wash buffer (0.1% SDS, 2mM EDTA, 20mM TrisHCl pH8, 1% TritonX-100, 500mM NaCl), once with LiCl wash buffer (0.25M LiCl, 1% IGEPAL-CA630, 1% deoxycolate, 1mM EDTA, 10mM TrisHCl pH8) and once with TE buffer (10 mM Tris-HCl (pH 7.5), 1mM EDTA). Chromatin was eluted by incubating the beads with elution buffer (1% SDS, 0.1M NaHCO3) for 15 min at RT. Eluted chromatin was reverse cross-linked by adding 10ng/ml of Proteinase K (Thermo Scientific) and 0.1M NaCL. After 2h incubation at 42°C, samples were incubated at 65°C O/N. Following morning, 10ng/ml of glycogen (Invitrogen) and 1 volume of Phenol:Chloroform:Isoamyl (25:24:1, Invitrogen, 15593031) were added and solutions were mixed thoroughly before isolating the DNA with Maxitract High Density Tubes (Qiagen).2.5 Volumes of 100% ethanol and 1/10 volume of 3M NaOAc (Sigma) were added to the sample and incubated for 15 min at −80°C. DNA was washed with 70% ethanol and resuspended in nuclease free water.

### ATAC-seq

ATAC-seq was performed as previously described^26^ with minor adaptations. Oli-neu cells (differentiated, siRNA treated as described above) were collected with Accutase solution (Sigma-Aldrich, A6964). 50 000 cells per condition were lysed in lysis buffer (10 mM Tris-HCl, pH 7.4, 10 mM NaCl, 3 mM MgCl2 and 0.1% IGEPAL CA-630) and centrifuged at 500xg at 4°C for 20 minutes. The cells were then resuspended in tagmentation mix (2.5 µl TD buffer, 2.5 µl Tn5 enzyme, Illumina) and the DNA was transposed for 30 minutes at 37°C, where after the DNA was purified using the Qiagen MinElute kit. After PCR amplification with 7-8 cycles (cycles determined with qPCR) the libraries were sequenced on a HiSeq2500 Illumina sequencer. Three replicates per condition were performed in different days and different passages. Each replicate had > 10 million reads with an average of 45 million reads per sample.

### Bioinformatic analysis of ChIP-seq and ATAC-seq

#### siPadi2 ATAC-seq processing

ATAC-seq siCtrl and si*Padi2* replicates were processed separately following the standard ENCODE pipeline for ATAC-seq samples, https://www.encodeproject.org/pipelines/ENCPL792NWO/. Adapters were detected and trimmed with cutadapt 1.9.1^46^ and aligned to mouse genome (GRCm38/mm10) using Bowtie2^47^, with default parameters. After filtering mitochondrial DNA, reads properly paired were retained and multimapped reads, with MAPQ < 30, were removed using SAMtools^48^. PCR duplicates were removed using MarkDuplicates (Picard - latest version 1.126), http://broadinstitute.github.io/picard/.

ATAC-seq peaks were called using MACS2 https://github.com/taoliu/MACS/^49^, with parameters -g mm -q 0.05 –nomodel –shift -100 –extsize 200 -B –broad. To get the consensus promoter/enhancer regions from each of the replicates we first selected the significant peaks −log10(p-value) > 1.30103. Using bedtools.2.17 mergeBed, we recovered the consensus ATAC peak region that included more than one replicate for siCtrl samples and si*Padi2* samples. From where we recovered 76021 and 55772 union peaks regions for siCtrl and si*Padi2*, respectively. For visualization and further analyses, the different replicates were merged using Samtools and tracks were normalized using deeptools bamcoverage, normalized with RPKM.

#### H3K27ac ChIP-seq data

Raw reads were mapped to the mouse genome (GRCm38/mm10) with Bowtie 2.1.0^47^. Uniquely mapped reads (17 and 19.5 million for proliferation and differentiation chromatin samples) were then processed with SAMTools for format conversion and removal of PCR duplicates^48^. Broad peaks corresponding to H3K27ac enrichment were called using MACS 2.0.10, with q-value cutoff at 5*10-249. H3K27ac peaks were filtered against blacklisted genomic regions prone to artifacts (ENCODE Project Consortium, 2012) with BEDTools^50^ and annotated to the nearest Transcription Start Site with PeakAnalyzer^51^.

#### Defined promoter/enhancer regions

We recovered the regions around the transcription start site (TSS) from gene annotations, ENSEMBL83. The putative promoter regions were extracted as -3KB upstream the TSS and 100bp down-tream the TSS, to recover possible promoter and enhancer regions. To get more reliable signal we intersected the siCtrl union peaks with peaks from H3K27ac ChIP-Seq in Olineu cells in differentiation and proliferation conditions (see above), ending with 17281 siCtrl/promoter peaks overlapping with H3K27ac peaks. Only the union peaks (7087 peaks) from siCtrl/H3K27ac overlapping the defined promoter regions were retained, using Bedtools intersect^50^. In order to recover possible enhancer and promoter regions that could be have been excluded from H3K27ac filtering or due to been more downstream of defined promoter regions, we included in the analysis a known mark of promoters/enhancers, H3K4me3 in OPC-like cells (GSE80089)^27^. We recovered the peaks bed file of H3K4me3 control peaks, GSM2112628, (liftover mm9 to mm10) and with bed-tools intersect, we recovered 10428 peaks corresponding to siCtrl/promoter/enhancer peaks with H3K4me3 peaks regions. The final dataset of peaks included the union of both promoter/enhancer peaks marked by H3K27ac and/or H3K4me3, with a total of 10897 peaks. The final region was extended 200bp up and downstream for further calculations. For the differential accessibility analysis between conditions the merged replicates from both conditions were used with Pyicos^52^ on the final peaks dataset, with parameters pyicoenrich –tmm-norm and default parameters. Including the z-score associated p-value and Benjamini & Hochberg (1995) corrected p-value (adj p-value or FDR)^53^ and the log2 fold change for candidates selection, with adjust p-value < 0.05 significance threshold. Data was visualized with IGV, http://www.broadinstitute.org/igv.

#### Gene ontology analysis

For the GO analyses, the top 300 up genes from siCtrl selected peaks and top 300 genes form si*Padi2* selected peaks, based on log2(RPKM siCtrl / RPKM si*Padi2*) were selected (adjusted p-val <0.05). GO and pathway analysis was performed with the ClueGO (version 2.5.1)^54^ plug-in Cytoscape (version 3.5.1)^55^ with settings, GO Biological process (20.11.2017) and REACTOME pathways (20.11.2017), showing only pathways with p-val <= 0.05. Default settings were used and a minimum of 4 genes per cluster were used.

### Statistical analysis

Statistical analysis was performed in GraphPad Prism7. Parametric and non-paired t-test was used to determine statistically significant differences between 2 groups in qRT-PCR experiments at the exception of the rat primary cultures where a Wilcoxon matched paired test was performed. One way ANOVA was used to determine differences between three groups in the behavioral analysis of *Padi2* fKO. Mann-Whitney tests were performed to compare Ct and *Padi2* cKO behavior.

### Accession numbers

The accession number for all raw data has been deposited in GEO GSE115929.

## Supplementary Materials

**Supplementary Table 1:** Citrullination targets detected in Oli-neu cells in both proliferation and 2-days differentiation. SILAC was performed in control and *Padi2* overexpressing Oli-neu cells with light peptides belonging to control and heavy peptides to *Padi2* overexpression cell lines.

**Supplementary Table 2:** PADI2 interactors detected in Oli-neu cells in both proliferation and 2-days differentiation. SILAC was performed in control and biotin tagged *Padi2* overexpressing Oli-neu cells. Light peptides belong to control and heavy peptides to biotin tagged *Padi2* overexpression cell lines.

**Supplementary Table 3:** Differential accessibility on siCtrl vs. si*Padi2* ATAC-seq peaks intersecting H3K4me3/H3k27ac marked promoters/enhancers.

**Supplementary Table 4:** Gene Ontology (GO) analysis for the citrullination targets, PADI2 interactors and differential accessible gene promoter/enhancer regions.

**Fig. S1.**
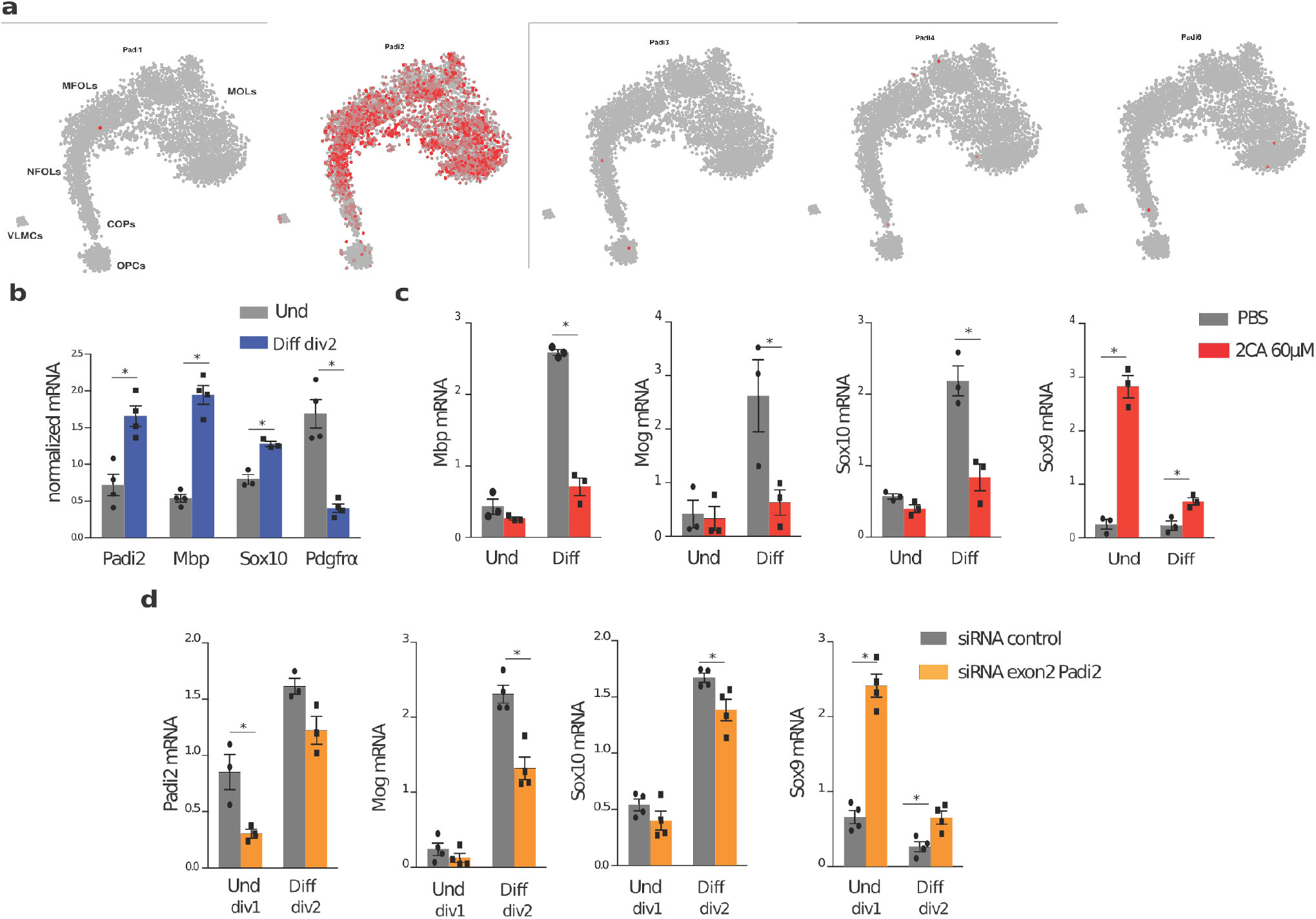
Inhibition of PADIs and *Padi2* knockdown impairs Oli-neu differentiation. (A) Gene expression of *Padi* family in the juvenile and adult OL lineage. Overlay in t-SNE from Marques *et al*.^15^. Scale gray - low expression, red - high expression). (B) Comparative gene expression analysis of undifferentiated (Und) and 2-days differentiated (Diff) Oli-neu cells. n=4, two-tailed t-test * p<0.05, error bars represent SEM. (C) Comparative gene expression analysis of undifferentiated (Und) and 2-days differentiated (Diff) Oli-neu cells treated either with PBS or with the PADI inhibitor 2CA. n=3, two-tailed t-test *p<0.05, error bars represent SEM. (D) Comparative gene expression analysis on undifferentiated (Und) and 1-day differentiated Oli-neu cells (Diff) transfected with either scramble siRNA or *Padi2* siRNA targeting exon2. Cells were collected at day in vitro (div)1 or div2 upon transfection. n=4, two-tailed t-test *p<0.05, error bars represent SEM.

**Fig. S2.**
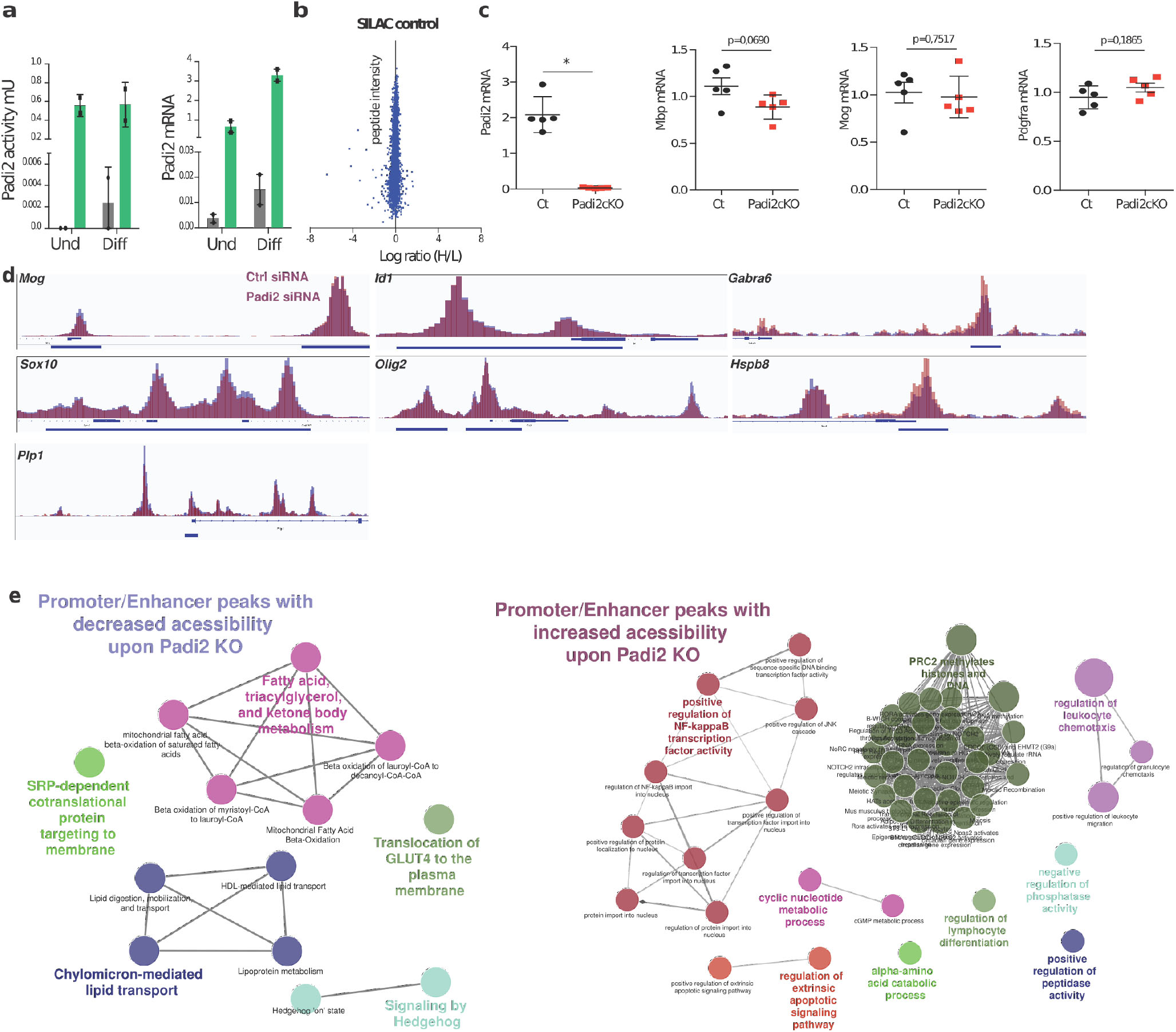
*Padi2* overexpression cell lines exhibit higher citrullination activity and modulation of chromatin accessibility upon *Padi2* knockdown. (A) Analysis of Oli-neu cell line overexpressing *Padi2* (green) or scramble control (black). *Padi2* overexpressing cells display higher PAD activity, as assessed by the ABAP kit assay and high mRNA *Padi2*, as assessed by qRT-PCR. (B) Incorporation of isotopes in Oli-neu cells was highly efficient once mixing proteins of LIGHT and HEAVY labeled control cells, resulting in an average ratio of 1, or log ratio (H/L)=0. (C) GFP+ cells were FACS sorted from controls (Ctr, *PdgfraCre*; RCE-loxP; *Padi2*+/+) and *Padi2* conditional knockouts (*Padi2* cKO, *PdgfraCre*; RCE-loxP; *Padi2*-/-). *Padi2*, *Mbp*, *Mog* and *Pdgfra* mRNA expression levels were compared. n=5, parametric t-test *p<0.05, error bars represent SEM. (D) IGV genome browser overlay views depicting chromatin accessibility (assessed by ATAC-Seq) near transcription start sites, in Oli-neu transfected with Ctrl siRNAs (blue) and siRNA against *Padi2*(red) (n=3, samples pooled for visualization, same scale for ctrl siRNA and *Padi2* siRNA). Regulatory regions (promoters/enhancers marked with H3K27ac or H3K4me3 in Oli-neu or OPCs, see methods) are highlighted with blue horizontal bars. (E) Gene ontology analysis of genes who present regulatory regions with increased or decreased chromatin accessibility upon *Padi2* knockdown in Oli-neu cells.

